# Non-Canonical RNA Binding of Human La-related Protein 6

**DOI:** 10.1101/2025.05.22.652967

**Authors:** Blaine H. Gordon, Victoria S. Ogunkunle, Robert Silvers

## Abstract

La-related proteins (LARPs) are RNA-binding proteins that are involved in a variety of disease-related processes. Most LARPs recognize short single-stranded poly(U/A) motifs via a conserved hydrophobic pocket. Human LARP6 (HsLARP6) is an exception, binding a structured 5′ stem-loop (5′SL) that controls type I collagen translation and fibroproliferative disease progression. Here, we present the *de novo* solution NMR structure of the La domain of HsLARP6 in the bound state. Chemical shift perturbation, solvent paramagnetic relaxation enhancement, intermolecular NOEs and targeted mutagenesis converge on a previously unknown binding interface that integrates electrostatic and hydrophobic contacts with shape complementarity in 5′SL binding. This non-canonical interface enables the La domain to discriminate 5′SL RNA from homopolymeric or purely helical hairpin RNAs with low-nanomolar affinity, overturning earlier views that the adjacent RRM is required for recognition. The structure provides the first molecular model for 5′SL recognition and expands the paradigm of La-mediated RNA binding beyond 3′-terminal oligo-U/A motifs. These insights provide the biophysical framework for molecular recognition of 5’SL by LARP6 that is related to collagen biosynthesis in fibrosis and associated pathologies.

## INTRODUCTION

La-related proteins (LARPs) constitute a superfamily of RNA-binding proteins (RBPs) that are involved in a variety of RNA-related processes such as RNA stability, metabolism, cellular localization, and gene regulation.[1–5] Due to their fundamental role in these cellular processes, LARPs are also associated with a variety of pathologies such as cancer,[6–14] viral infection,[15–19] auto-immune disorders,[20,21] and fibroproliferative disease.[22,23] Most LARPs target single-stranded RNAs depending on their function, most commonly the 3’ termini of poly(A) or poly(U) sequences of their target RNAs.[24–30] Human La-related protein 6 (HsLARP6), however, recognizes an RNA that is more complex in sequence and structure, known as the 5’stem-loop (5’SL).[31,32] The secondary structure of 5’SL RNA is predicted to contain an internal loop or bulge formed between two helical stems whose sequence spans the junction between 5’ untranslated and coding regions of type I and type III collagen mRNAs (Figure 1A, left). HsLARP6 modulates fibrillar collagen biosynthesis [33–40] and fibroproliferative disease progression [7,8,22,38,41–43] through associations with the 5’SL motif.[31,34,40,44–46] Through the localization and translation factors HsLARP6 recruits, these mRNAs are efficiently translated to produce properly modified, folded, and excretable collagen at rates several hundred-fold greater than those of constitutive collagen synthesis.[34–36,44,46–48]. *In vitro* mutation and binding studies of 5’SL sequence elements highlighted a lack of sequence specificity in stem regions, which are required to form the internal loop.[33,49] In contrast, single nucleotide mutations in five of the nine bulge nucleotides abolished binding entirely.[33,49,50]

**Figure 1.**
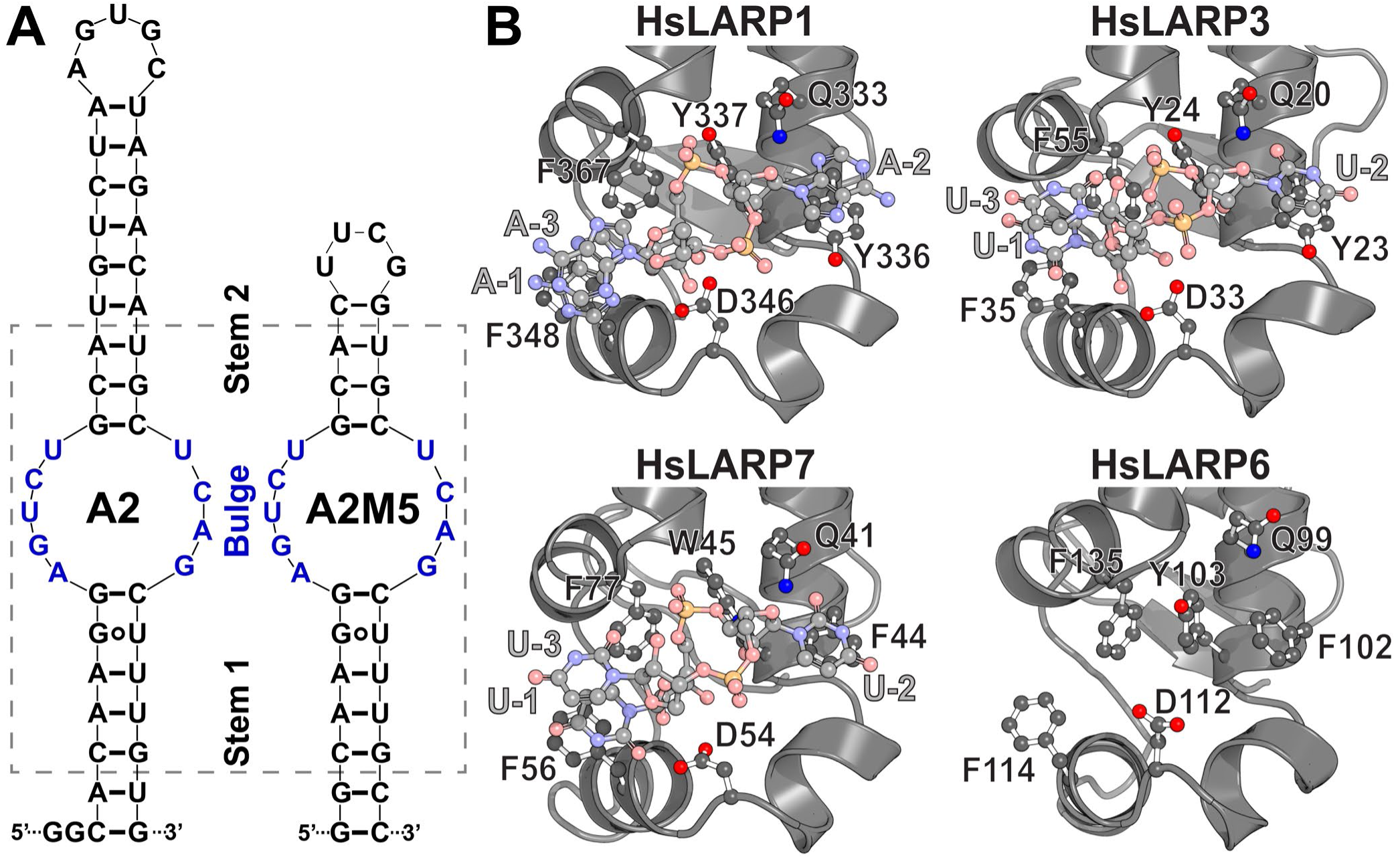
Canonical and non-canonical RNA binding of LARPs. (A) Sequences and secondary structure predictions of wild type A2 RNA (left) and the 35-mer A2M5 model RNA (right). Nucleotides shared between the wild type and model sequences are indicated by a gray, dashed box. (B) Close-up views of the canonical RNA binding pocket in structural models of La domains from HsLARP1 binding an AAA- 3’OH RNA (top left), HsLARP3 (top right) and HsLARP7 (bottom left) binding UUU-3’OH RNA are shown alongside the protein structure of the A2M5-bound state of the La domain of HsLARP6 (bottom right). Side chains of residues relevant for RNA binding are shown and labeled. Structural representations of HsLARP1, HsLARP3, HsLARP6, and HsLARP7 were generated using PDB entries 7SOR, 2VOD, 9NGX, and 4WKR, respectively.[28,30,53]

Canonical RNA recognition of single-stranded poly(A) or poly(U) sequences by HsLARPs 1, 3, and 7 was previously investigated by crystallographic and mutational studies. Generally, RNA binding in LARPs involves a highly conserved winged helix domain known as the La domain (Figure 1B) that often forms a bipartite RNA binding module with a downstream ribonucleotide recognition motif (RRM), named the La module.[25,30] However, deviation from this tandem domain model exists in HsLARP1 and HsLARP6. HsLARP1 was shown to lack a folded RRM domain in NMR spectroscopic studies of the La module, having only a La domain which binds poly(A) with low nanomolar affinity *in vitro*.[28] While the La module of HsLARP6 contains an RRM domain, it is not explicitly required for high affinity 5’SL binding.[50–52] X-Ray crystallographic structures of RNA-bound La domains and La modules of HsLARP1, HsLARP3, and HsLARP7 (Figure 1B) present a striking similarity in binding mode.[25,28,30] In all cases, the UUU-3’OH or AAA-3’OH RNA form a splayed-nucleotide orientation.

Canonical RNA binding in LARPs is achieved through a set of conserved amino acids performing virtually identical roles across all known La domain structures (Figure 1B, Figure S1).[28,30,53] The bases of nucleotides in position -1 and -2 are splayed apart and interact with conserved aromatic residues of helix α1 and α2 via π-stacking, respectively. The base of the nucleotide in position -3 stacks with the base in position -1 forming a π-stacking triple. A glutamine residue in helix α1 forms a hydrogen bond with base -2 while a tyrosine (tryptophan in HsLARP7) residue in helix α1 forms a hydrogen bond to a backbone phosphoryl oxygen located between bases -1 and -2. An aspartate in helix α1’ forms hydrogen bonds to the 2’- and 3’-hydroxyl moieties of the ribose of nucleotide -1, however, mutagenesis of this residue had only a small effect on RNA binding in HsLARP3. Aside from the aforementioned aspartate, mutation of these residues was shown to drastically diminish RNA binding.[25,28,54] These residues are conserved in HsLARP6 as well (Figure 1B, bottom right), despite 5’SL lacking a 3’OH motif found for target RNA of other HsLARPs. Hence, the molecular basis for 5’SL binding by the La domain of HsLARP6 remains unclear.

Here, we present the *de novo* solution NMR structure of the 5’SL-bound La domain of HsLARP6 and showed that the La domain employs a previously unknow, non-canonical binding interface to target 5’SL RNA. Chemical shift perturbation analysis, solvent PREs, intermolecular NOEs, as well as mutagenesis all point towards a distinctly different binding site for 5’SL unique to HsLARP6. Furthermore, we demonstrate the ability of the HsLARP6 La domain to discriminate between 5’SL RNA, homopolymer sequences, and purely helical hairpin RNAs. This work presents the first atomic-resolution model describing the molecular underpinning of the interaction between the HsLARP6 La domain and the 5’SL motif found in type I collagen mRNA.

## MATERIALS AND METHODS

### Sample preparation

Samples for NMR spectroscopy and MST were prepared as described previously.[55,56] In the following, we will briefly outline the procedures of protein and RNA production and purification. La domain constructs including cysteine and alanine point mutations cloned into a pET28a vectors via *Nco*I and *Bam*HI restriction sites for expression in *E. coli* were purchased from Genscript USA Inc. All construct names and protein sequences are listed in Table S1. Protein expression and purification proceeded as previously reported.[55,56] RNA was synthesized by *in vitro* transcription (IVT) as described previously,[55,56] unless stated otherwise. RNase-free and HPLC-purified template DNA and T7 promoter oligonucleotides used for IVT were obtained from Integrated DNA Technologies, USA. Sequences of DNA oligonucleotides are listed in Table S2. RNase-free and HPLC-purified RNA oligonucleotides of the 6-mer A6 and U6 as well as hairpin RNAs 14-mer (UUCG) and 14-mer (GAAA) were purchased from Genscript, USA. Sequences of RNA oligonucleotides are listed in Table S3.

### NMR Spectroscopy

All NMR experiments were acquired at 298K on a Bruker AVANCE III NMR spectrometer equipped with a TCI cryoprobe operating at 16.4 T operating TopSpin 3.6.5 unless stated otherwise. Many of the NMR experiments used here were previously used for resonance assignment and were published previously.[56] All NMR experiments were conducted on samples of the 1:1 complex of HsLARP6(79-183) and A2M5 RNA, unless otherwise stated. HsLARP6(79-183) was either uniformly [^15^N]-labeled or [^13^C, ^15^N]-labeled while all RNA was natural abundance (NA). NMR samples were measured in 10 mM MES pH 6.5, 50 mM KCl, 10% D_2_O or 10 mM MES pD 6.5, 50 mM KCl, 100% D_2_O. DSS was added to all samples at a final concentration of 0.01 mg/mL as an internal ^1^H chemical shift reference.[57] ^13^C and ^15^N resonances were referenced indirectly.[58] Data was processed with TopSpin 4.2.0 and analyzed using NMRFAM-SPARKY 1.47.[59,60]

### Resonance assignment of unbound and bound La domains

The near-complete ^1^H, ^13^C, and ^15^N resonance assignment of the 1:1 complex of the La domain of HsLARP6 and A2M5 RNA was published previously (BMRB accession code: 31231).[56] Briefly, a standard suite of triple resonance experiments including 3D HNCA, 3D HNCACB, 3D HNCO, and 3D HN(CA)CO were used for backbone assignments,[61–70] while side chain assignments heavily relied on 3D (H)CC(CO)NH, 3D H(CC)(CO)NH, 3D HCCH-TOCSY, 3D (H)CCH-TOCSY, and 3D HCCH-COSY spectra.[71–78] Additionally, assignments were aided by a series of ^15^N- and ^13^C-edited 3D NOESY-HSQC experiments described below that were also used for structure determination (*vide infra*).[79,80] Detailed descriptions of NMR samples as well as data acquisition and processing of these spectra can be found in Gordon and Silvers (2025).[56]

Backbone amide assignment of the unbound state of HsLARP6(79-183) was performed by comparison of the [^1^H, ^15^N]-HSQC spectrum to the previously published assignment of the unbound state of HsLARP6(70-183) deposited in the BMRB (BMRB ID 25159).[52] This was necessary due to the substantially different buffer conditions and the difference in length at the N-terminus. Backbone amide assignment of the A2-bound state of HsLARP6(79-183) was performed by comparison of the [^1^H, ^15^N]- HSQC spectrum to the previously published assignment of the A2M5-bound state of HsLARP6(79-183) deposited in the BMRB (BMRB ID 31231).[56] [^15^N]-HSQC spectra of 500 μM [^15^N]-labeled unbound HsLARP6(79-183) (Figure S2) and 200 μM [^15^N]-labeled A2-bound HsLARP6(79-183) (Figure S3) in buffer containing 10 mM MES pH 6.5, 50 mM KCl, 10% D_2_O, and 0.01 mg/mL DSS were recorded with 2048 x 256 points. Backbone amide assignments of the unbound and A2-bound state of HsLARP6(79-183) can be found in Tables S4 and S5, respectively.

### Structure determination of the A2M5-bound La domain

Distance restraints derived from a set of six individual ^15^N- and ^13^C-edited 3D NOESY-HSQC spectra were acquired for use in structure determination. One ^15^N-edited and ^13^C-edited 3D NOESY-HSQC spectrum each were acquired on a uniformly [^13^C, ^15^N]-labeled sample. During processing the ^13^C-edited 3D NOESY-HSQC spectrum was split into two separate spectra (aliphatic and aromatic regions) for easier analysis. Additionally, individual aliphatic and aromatic ^13^C-edited 3D NOESY-HSQC spectra of [^15^N, 2-^13^C-glycerol]-labeled and [^15^N, 1,3-^13^C-glycerol]-labeled samples were acquired. The use of alternating carbon-13 labeling introduced by the 1,3-^13^C-glycerol and 2-^13^C-glycerol to aid in resonance assignment has been previously described,[56,81] and, similarly, the overall decrease in congestion and linewidth in strongly coupled aromatic side chains aided in the extraction of distance restraints.

Peaks in all 3D NOESY-HSQC spectra were picked manually using NMRFAM-SPARKY, except for the ^13^C-edited 3D NOESY-HSQC spectra that were split into the aliphatic and aromatic regions. For these 2 spectra, automatic peak picking was performed using the ARTINA[82] peak picking algorithm on the NMRTIST[83] platform followed by manual curation in NMRFAM-SPARKY to remove picked peaks belonging to artifacts at the edges of the spectrum and close to the water. Prediction of torsional angles was performed using TALOS-N.[84]

Resonance assignments, dihedral constraints derived from TALOS-N[84], and the peak lists derived from manual and automatic peak picking were subjected to the combined automated NOE assignment and structure calculation routine implemented in CYANA 3.98.15 using the *noeassign* routine as described previously.[85–87] The commands *ramaaco* and *rotameraco* were used to restrict backbone and side chain dihedral angles, respectively, to the most favored regions. 20 structures with the lowest target function out of 400 were selected and subjected to refinement using AMBER 22[88,89].

### Dynamics of the A2M5-bound La domain

^15^N longitudinal (R_1_) and transverse (R_2_) relaxation rates, as well as (^1^H)-^15^N heteronuclear steady-state NOEs (hetNOEs) were determined using established protocols.[90,91] All relaxation experiments were recorded on a sample containing 400 μM of the 1:1 complex of [^15^N]-HsLARP6(79-183) and [NA]-A2M5 in 10 mM MES pH 6.5, 50 mM KCl, 10% D_2_O, and 0.01 mg/mL DSS. The R_1_ relaxation experiment was recorded with 12 relaxation delays of 0.02, (2x) 0.1, 0.2, (2x) 0.4, 0.6, 0.8, (2x) 1.2, 1.6, 2.0 s. The R_2_ relaxation experiment was recorded with 12 relaxation delays of 17, (2x) 34, 68, (2x) 102, 136, 170, (2x) 204, 238, and 272 ms. The hetNOE pseudo-3D was recorded with a recycle delay of 6.0 s. Experiments were recorded as pseudo-3Ds and after processing, individual 2D planes were extracted in TopSpin and intensities were subsequently analyzed using NMRFAM-SPARKY[60]. Reduced spectral density mapping was performed using RELAX 5.1.0 as described previously.[92–96] The molecular weight of the A2M5-bound complex was determined by using a strategy described previously (https://nesgwiki.chem.buffalo.edu/index.php/NMR_determined_Rotational_correlation_time).

### Chemical Shift perturbations and solvent PREs

Using the resonance assignments of the unbound, A2M5-bound, and A2-bound La domain (*vide supra*), the pairwise chemical shift perturbations (CSPs) for each residue were determined using the weighted Euclidean distance between peak with a scaling factor (α) of 0.1 as described previously.[97] To determine the protein surface protected from paramagnetic relaxation enhancement by the binding of 5’SL, [^1^H, ^15^N]-HSQC spectra of a 200 μM [^15^N]-labeled A2-bound HsLARP6(79-183) sample in buffer containing 10 mM MES pH 6.5, 50 mM KCl, 10% D_2_O, and 0.01 mg/mL DSS were recorded with 2048 x 256 points. From a 50 mM stock solution of gadodiamide (Gd-DTPA-BMA) in the same buffer, the NMR sample was titrated to a final concentration of 4, 8, and 12 mM gadodiamide. For each concentration, a [^1^H, ^15^N]-HSQC spectrum was recorded. Signal attenuation (I/I_0_) at each concentration was calculated by taking the ratio of the signal intensities from each titration point (I) versus the reference spectrum with no added gadodiamide (I_0_).

### Intermolecular contacts between La domain and A2M5

Intermolecular NOEs were identified between the imino and methyl regions of a 2D [^1^H, ^1^H]-NOESY of a sample containing the 1:1 complex of 630 μM natural abundance HsLARP6(79-183) and natural abundance A2M5 with 3072 x 768 points, water suppression using 1-1 echo pulses sequence, and a mixing time of 250 ms.[98] Additionally, a 2D ^15^N-edited [^1^H,^1^H]-NOESY-HSQC spectrum a sample containing the 1:1 complex of 1.9 mM [^13^C,^15^N-Arg]-labeled HsLARP6(79-183) and [NA]-A2M5 was recorded with 1664 x 640 points and a mixing time of 200 ms.

### Microscale Thermophoresis

Binding affinities of RNA binding to the La domain of HsLARP6 were determined by employing microscale thermophoresis (MST) as previously described.[55,56,99] Briefly, Cy5 is attached to LARP6(73-183) via a cysteine mutation (G74C) present in the flexible N-terminus of the La domain. We termed this Cy5-labeled “wild type” protein Cy5-HsLARP6(73-183), while also using this construct to determine binding affinities of single-site alanine mutations in the core domain. For a full list of constructs and sequences, see Table S1. Cy5-labeled protein and refolded RNA species were separately exchanged into a buffer containing 20 mM Tris pH 7.25, 100 mM KCl, 5 mM DTT, and 0.05% Triton X-100 by gel filtration on a Superdex 75 Increase 10/300 GL column (Cytiva, USA). Both protein and RNA were supplemented with final concentrations of 0.4 mg/mL BSA and 0.2 mg/mL yeast tRNA from their respective 20 mg/mL and 10 mg/mL stock solutions. Cy5-labeled protein was kept at a constant concentration of 10.0 nM in all MST measurements, except for the variant Cy5-HsLARP6(73-183)F102A where the concentration was 20.0 nM. Titration of RNA was achieved by 16 1:1 serial dilutions. For A2M5 RNA the final concentrations in the capillaries ranged from 5 μM to 152.6 pM for all variants, except for the variant Cy5-HsLARP6(73-183)F114A where the final concentrations in the capillaries ranged from 20 μM to 610.4 pM. For hairpin RNAs HP1 and HP2 the final concentrations in the capillaries ranged from 5 μM to 152.6 pM. For the hairpin RNAs 14-mer (UUCG) and 14-mer (GAAA), as well as the single stranded 6-mers A_6_ and U_6_, the final concentrations in the capillaries ranged from 100 μM to 3.05 nM.

The normalized fluorescence (F_NORM_) for each ligand concentration was determined by dividing the average fluorescence signal between 0.5 s to 1.5 s after the IR laser on-time and the average fluorescence signal between -1.0 s to 0.0 s before the IR laser on-time for each capillary. All measurements were collected with three replicates. For RNA showing binding, the fraction bound and apparent dissociation constants (K_d_) were determined from thermophoresis profiles using procedures described previously.[55,56] For non-binders, the change in normalized fluorescence (ΔF_NORM_) was determined by subtracting all data points from the data point at the lowest ligand concentration.

## RESULTS

### A2M5 as a model for 5’SL RNA

The 5’SL RNA found in type I collagen mRNA is around 46-48 nucleotides in length,[31] with extensive sections predicted to be base-paired and helical (Figure 1A, left). While the 5’SL-bound structure presented here is focused entirely on the protein of the complex, any future NMR spectroscopic studies that involve spectra of 5’SL RNA are limited by the presence of resonance overlap commonly present in RNA spectra. To reduce the molecular size of the complex and minimize spectral overlap in RNA-based spectra, a shorter 35-mer 5’SL model named A2M5 was designed (Figure 1A, right). Since 5’SL-binding is independent of the stem sequence and length,[49,50] A2M5 was designed with a significantly shortened stem 2, with the length of stem 1 mostly unaltered. The internal loop itself as well as nucleotides adjacent to it are unchanged (Figure 1A, box). We previously showed that A2M5 is an excellent model of reduced size with unchanged binding characteristics compared to the wildtype A2 sequence.[56] Binding studies using a model 14-mer RNA with a cUUCGg and a cGAAAg tetraloop show no affinity the La domain (*vide infra*, Figure S14), indicating that the presence of exposed nucleobases in the loop do not contribute substantially to binding. Comparison of [^1^H, ^15^N]-HSQC spectra and chemical shift perturbation analysis revealed only minor differences between the A2-bound and A2M5-bound state (Figures S3 and S4). Minor chemical shift perturbations are observed for the backbone amide resonances of K139 and H140, indicating that these residues residing in helix α4 are in close proximity to stem 2 adjacent to the UUCG loop. Notably, this provides an important clue as to the location of stem 2 and the general orientation of the 5’SL RNA when bound to the La domain.

### The structure of the 5’SL-bound La domain of HsLARP6

The 23.9 kDa complex formed between the La domain of HsLARP6 and the A2M5 model RNA was used for structural and biophysical investigation using solution-state NMR spectroscopy (Figure 1A, right). We recently reported on the near-complete backbone and side chain resonance assignment of the A2M5-bound La domain of HsLARP6 with a total of 99.3% of observable protons assigned (BMRB: 31231).[56]

Here, a data set consisting of distance restraints derived from six 3D NOESY-HSQC spectra combined with dihedral angle restraints were subjected to the combined automatic NOE assignment and structure calculation using CYANA (Table S6).[85] 20 structures with the lowest target function were selected and subsequently subjected to water refinement using Amber (Figure 2B).[88] 86.6% of backbone torsion angles fall within the most favored region of the Ramachandran plot, with the remaining 13.4% falling in the additionally allowed regions. The well-defined core structure of the La domain of HsLARP6 spanning residues W85 to F176 was determined with average backbone and heavy atom RMSDs of 0.21 ± 0.04 Å and 0.62 ± 0.05 Å, respectively. The N-terminal and C-terminal residues are less well-defined and flexible. The average backbone and heavy atom RMSDs of the full structure spanning residues G78 to S183 was 1.69 ± 0.34 Å and 1.76 ± 0.28 Å, respectively.

**Figure 2.**
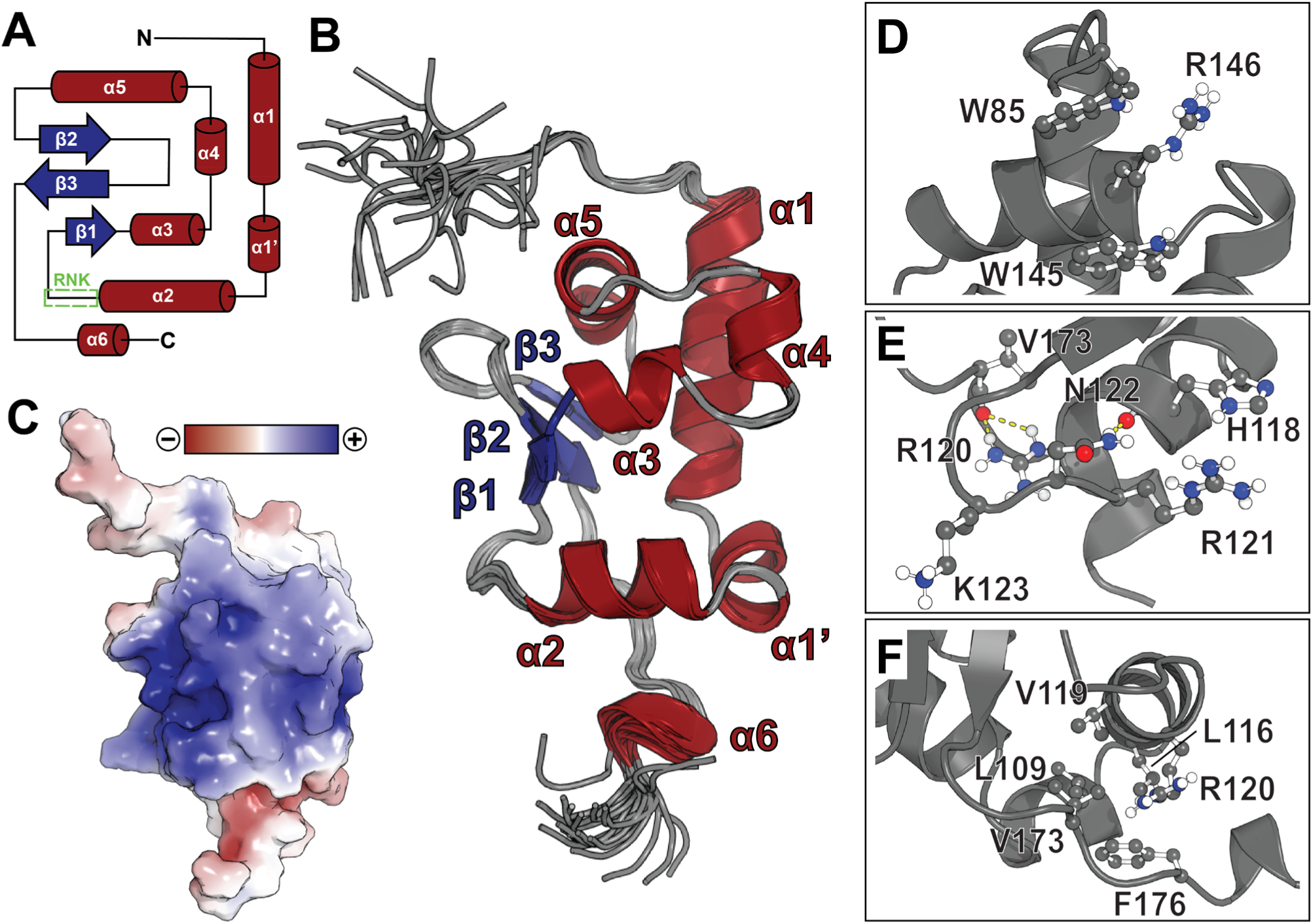
Structure and topology of the A2M5-bound La domain of HsLARP6. (A) Secondary structure topology of the La domain of HsLARP6. (B) The 20 structures of the A2M5-bound La domain of HsLARP6 as determined by the solution NMR spectroscopy (PDB: 9NGX). The structures are shown cartoon representation with α-helices, β-strands, and loops are shown in red, blue, and gray, respectively.(C) The surface charge of the La domain as determined by APBS shows a continuous basic patch (blue).[100] The structure is oriented identically to the structure shown in panel B. (D-F) Sections of interest show structural details surrounding residues W85/W145 (D), the RNK motif (E), and the hydrophobic cluster involving the C-terminus (F). Side chains of select residues are shown and labeled.

The structure of the 5’SL-bound La domain of HsLARP6 presents with a secondary structure topology consisting of six α-helices and three β-strands forming a winged helix motif known for La domains (Figure 2A),[1–5,25,27,28,101], and as was previously predicted by chemical shift analysis.[56] An additional 3_10_ helix (α6) is located at the C-terminus. Residue R146 located in helix α5 is positioned between W85 and W145, as evidenced by strong NOE contacts and chemical shifts of H^β2/3^ (1.054/0.373 ppm) and H^γ2/3^ (0.059/-0.152 ppm) nuclei of R146 experiencing strong upfield shifts due to the ring currents of both close-proximity tryptophan residues (Figure 2D). The region known as the RNK motif spanning residues R121, N122, and K123 is well-structured with the side chain of N122 forming hydrogen bonds to the C=O moiety of residue H118 (Figure 2E). Likewise, the side chain of R120 forms hydrogen bonds with C=O of residue V173. The hydrophobic residues located at the C-terminus, in particular residues L109, L116, V119, V173, and F176 form a hydrophobic cluster that includes the aliphatic side chain of R120 (Figure 2F). Surface residues that are located in helices α2, α3, α4, α5, the RNK motif, and strand β1 form a large basic patch due to the presence of many lysine, arginine, and histidine residues (Figure 2C). Interestingly, this basic patch is a unique feature of HsLARP6 (Figure S5) and contributes to nucleic acid binding via salt bridge formation between the protein and the negatively charged phosphate backbone of the RNA, a common theme in nucleic acid binding.[102–106]

### The unbound and 5’SL-bound states of the La domain differ substantially in structure and dynamics

Alignment of the unbound (PDB: 2MTF)[51] and A2M5-bound structures of the La domain revealed significant r.m.s deviations (RMSDs) within the well-defined core structure of the La domain (Figure 3). Most notably, the RNK motif (residues 121-123) was more rigid in the presence of the bound RNA. In the unbound state, the amide resonances of many residues located in the RNK motif were unobserved due to conformational exchange,[51,52] but were present and assigned in the bound form (*vide infra*). This is also supported by the observation that the μs-ms dynamics within the RNK motif are quenched upon binding of RNA. Furthermore, the region surrounding residue W145 and the loop located between helices α4 and α5 is shifted towards the N-terminal end of helix α3 with a noticeable change in angle of helix α5. Overall, it is noticeable that the bound structure converges on a more uniform model than the unbound state. The A2M5-bound state of the protein forms a continuous hydrophobic core that is not present in the unbound state, likely contributing to the change in protein stability upon RNA binding as was previously described.[55]

**Figure 3.**
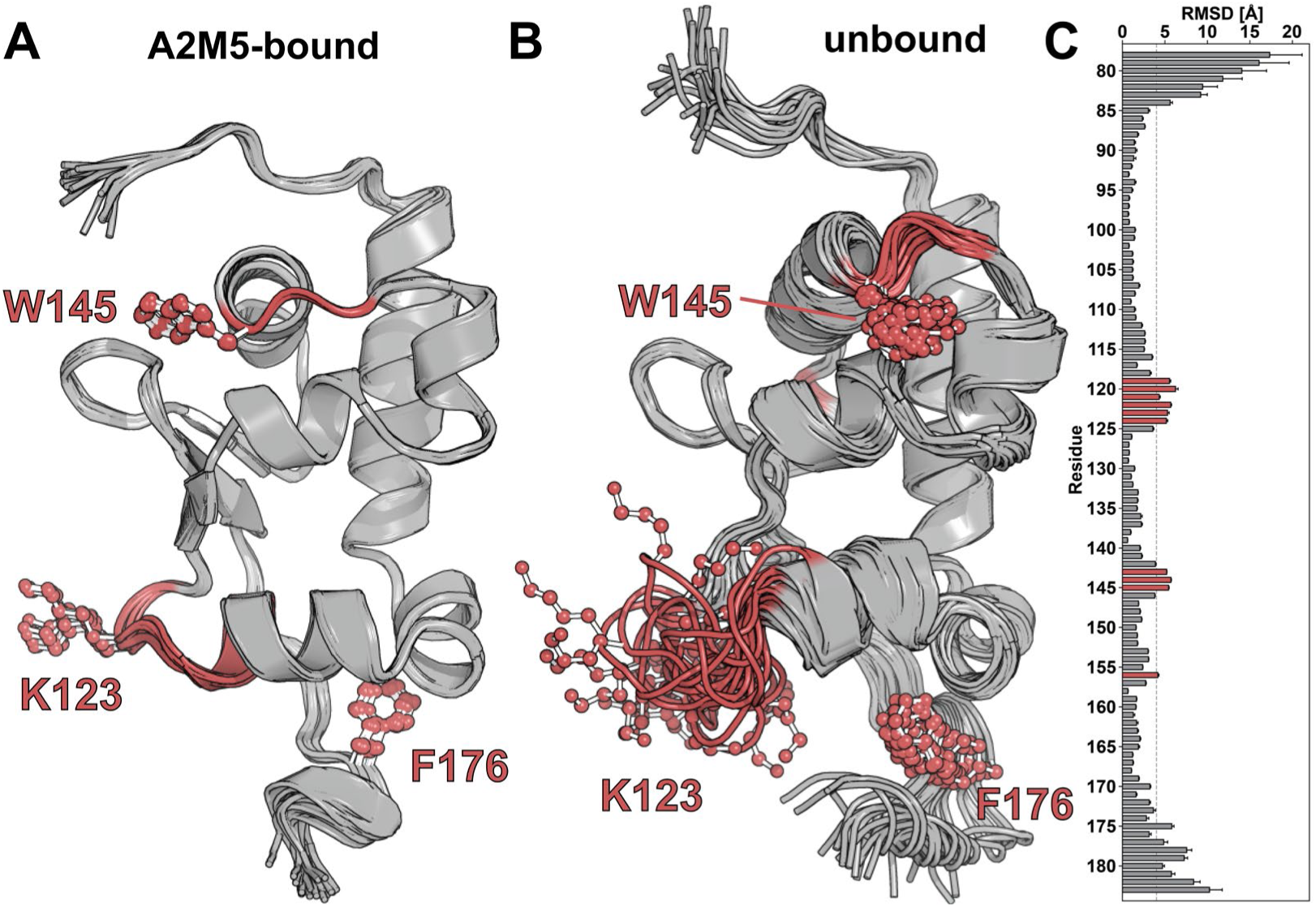
Comparison of the A2M5-bound and unbound structures of the La domain of HsLARP6. The structures of the A2M5-bound La domain (A, PDB: 9NGX) and unbound La domain (B, PDB: 2MTF) are shown in cartoon representations. Regions with RMSD values above 4.0 Å are shown in red and side chains are shown for select residues. (C) Average residue-specific root mean square deviations (RMSDs) between the structural ensembles of the A2M5-bound and unbound structures. RMSD values above 4.0 Å within the well-defined core structure excluding the more flexible N- and C-terminal tails are shown in red.

Backbone dynamics on the pico-to-nanosecond timescale were probed using longitudinal (R_1_) and transverse (R_2_) ^15^N relaxation rates as well as steady-state ^1^H-^15^N hetNOEs (Figure S6, Table S7).[90] Rotational correlation times of the core domain as determined from T_1_/T_2_ ratios were used to determine an approximate molecular weight of 25.0 ± 1.3 kDa, in good agreement with the expected molecular weight of the 1:1 complex of 23.9 kDa (Figure S7). Reduced spectral density mapping revealed that residues showing chemical exchange contributions are either located in the vicinity of the RNK motif (L124 and V129) or within helix α4 (K139 and H140), likely due to experiencing conformational exchange on the μs-ms timescale (Figure S8).[92,93] Additionally, the amide resonance of residue K138 is not observed due to exchange broadening.[56]

### The non-canonical binding interface of the La domain of HsLARP6

The amino acid residues of LARP6 that are directly involved in 5’SL binding are mostly undetermined. This is largely due to the focus on residues in the La module of HsLARP6 analogous to those of the canonical binding site in other HsLARPs. It has been previously proposed that the La domain of HsLARP6 utilizes the same conserved residues to recognize 5’SL RNA based on observations from the binding of single-stranded RNA of other LARPs.[51] Recently, however, UV cross-linking experiments have shown that residues located in the RNK motif are involved in RNA binding, which are unique to LARP6 and conserved among vertebrates.[50]

First, to identify the interface employed by the La domain in 5’SL binding, we employed a combination of chemical shift perturbations (CSPs), solvent paramagnetic resonance enhancement (sPRE), and intermolecular NOEs (Figure 4). The change in chemical environment upon 5’SL binding was observed by CSP analysis using the ^1^H-^15^N amide moieties as probes (Figure 4A&D). Significant CSPs (> 0.1 ppm) were not observed within the canonical La domain binding site. Instead, 5’SL binding had the most pronounced effect on the chemical shift of residues surrounding residues L115, L132, and K139 (Figure 4A&D, dark red). Some CSPs larger than 0.1 ppm were identified for residues not located at the protein-RNA interface. These CSPs are related to allosteric effects in regions distant from the main binding site, which is in line with the observation that minor CSPs are observed throughout the structure and in particular for residues involved in forming the continuous hydrophobic cluster (*vide supra*).

**Figure 4.**
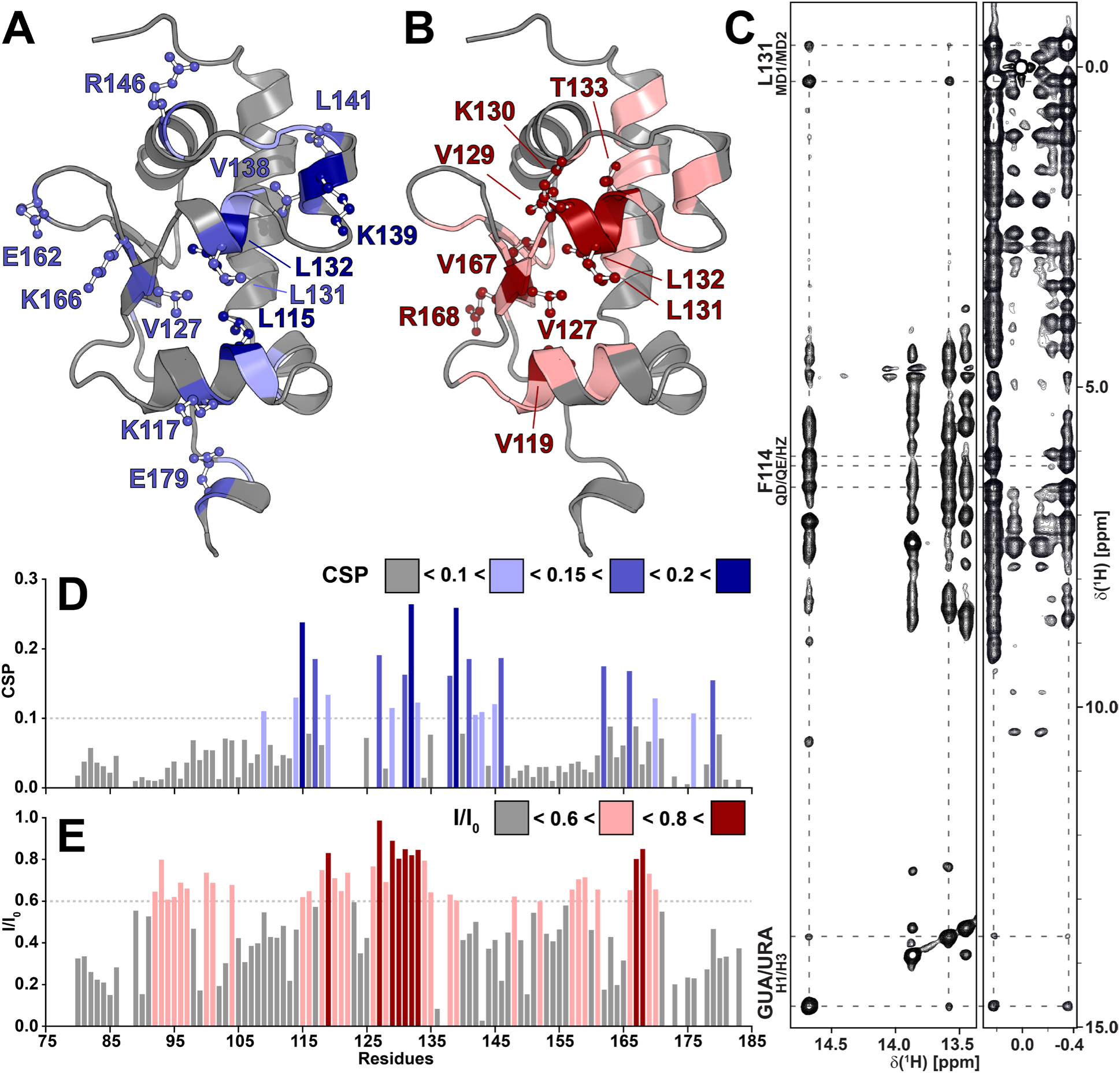
Mapping the binding surface for 5’SL RNA on the La domain of HsLARP6. (A) Chemical shift perturbations (CSPs) between the A2M5-bound and unbound La domain are shown on a representative model of the A2M5-bound state. Residues with CSPs below 0.1 ppm are colored gray, residues with CSPs between 0.1 and 0.15 ppm, between 0.15 and 0.2 ppm, and above 0.2 ppm are colored light blue, blue, and dark blue, respectively. Side chains are shown and are labeled for residues with a CSP > 0.15 ppm. (B) Solvent accessibility is shown on a representative model of the A2M5-bound state as reported by the ratio of NMR signal intensity in the presence (I) or absence (I_0_) of co-solvent gadodiamide. Highly (I/I_0_ > 0.8) and moderately (0.8 > I/I_0_ > 0.6) inaccessible residues are colored in dark and light red, respectively. Side chains are shown and are labeled for residues with an I/I_0_ > 0.8. (C) 2D ^1^H-^1^H-NOESY spectrum showing intermolecular NOE contacts between A2M5 RNA and the La domain. Parts of the imino region (left) and methyl region (right) are shown. Dashed lines link diagonal and cross peaks observed in both spectra while labels indicate the identity of the observed residues from each molecule. (D+E) Bar plots of CSP and I/I_0_ values. Dashed lines indicate the minimum threshold for categorization for each data set. Color-coding is as described for panels A and B.

Solvent accessibility was probed by NMR signal attenuation in backbone amides caused by titration of the paramagnetic cosolvent gadodiamide, a common MRI contrast agent (Figures 4B&E, S9). When gadodiamide was added to a sample of the 1:1 complex of [^15^N]-HsLARP6(79-183) and full-length A2 RNA at 60-fold excess, the residues with the least signal attenuation (I/I_0_ > 0.8) resided in helices α2 and α3, the RNK motif, and strand β1 (Figure 4B&E). These residues are well protected from the paramagnetic relaxation enhancement due to the binding of 5’SL RNA.

While a significant portion of proton chemical shifts in proteins and nucleic acids overlap, some intermolecular NOEs can be identified even in the absence of unambiguous resonance assignment of the A2M5 RNA. Particularly prominent are the cross peaks between the imino protons of two nucleobases (uracil or guanine) to the methyl groups of L131 and the close-by side chain of F114 (Figure 4C and S10).

Notably, both imino protons are involved in base-pairing and are in close proximity as evidenced by the cross-peak between both imino protons. Additionally, the ^15^N-edited 2D NOESY-HSQC spectrum of a sample of the 1:1 complex of [^13^C,^15^N-Arg]-HsLARP6(79-183) and natural abundance A2M5 showed cross-peaks between the side chain H^ε^ of R121 and several imino protons of A2M5 (Figure S11).

Taken together, chemical shift perturbations, solvent accessibility, and intermolecular NOEs are consistent, pointing towards a 5’SL binding surface involving helices α2, α3, α4, and the RNK motif. Notably, the binding interface matches the location of the basic patch (Figure 1C).

### Site-directed mutagenesis reveals key residues

Alanine screening was employed to identify residues directly involved in the binding of 5’SL RNA. We determined the binding affinities between A2M5 and 16 alanine point mutants of various aromatic and positively charged residues lining the binding interface of the La domain (Figures 5 and S12).

**Figure 5.**
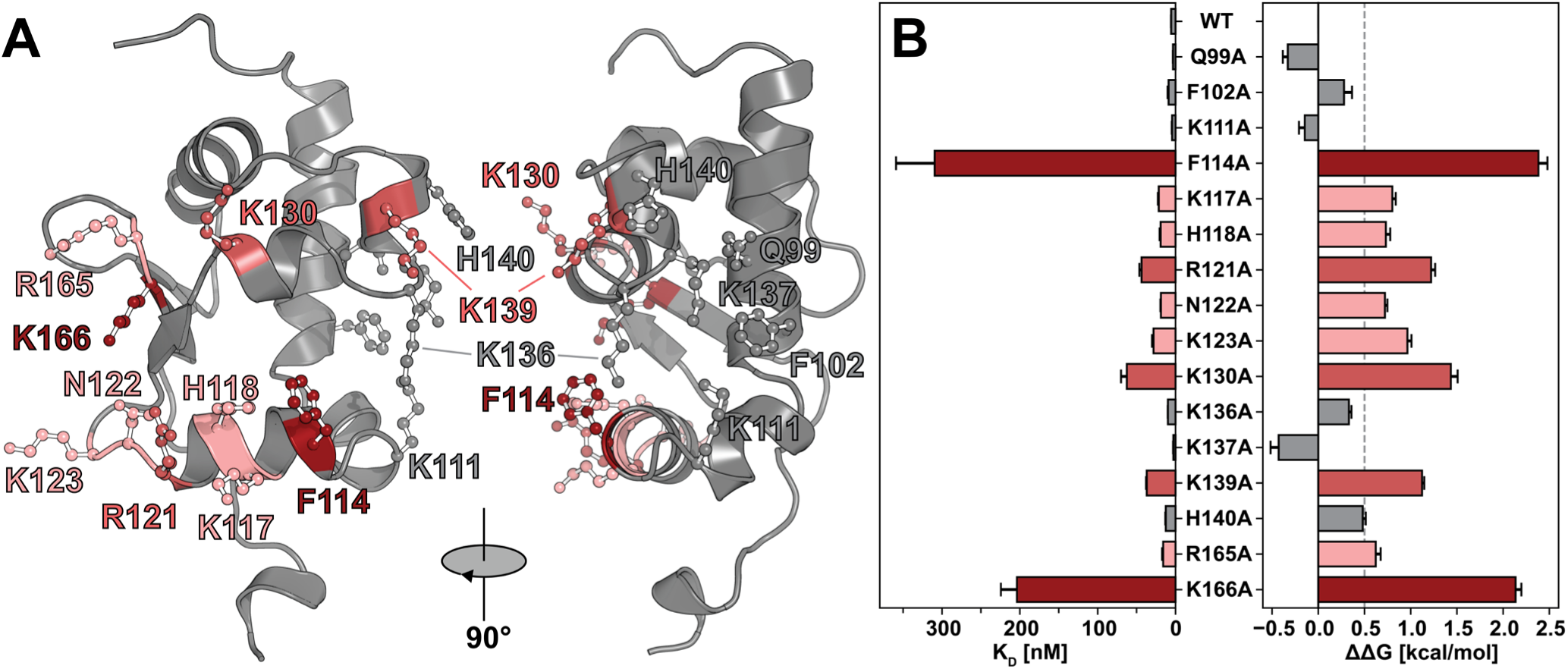
Mutagenesis of key residues in the La domain of HsLARP6. (A) Locations of single-point alanine mutants throughout the La domain are shown on a representative structure of the A2M5-bound structure. (B) K_D_ and ΔΔG values determined for each measurable mutation (Table S8). Mutated residues are color-coded based on the relative effect their mutation had on the dissociation constant (K_D_) of A2M5 binding. ΔΔG values above 2.0 kcal/mol are shown in dark red, ΔΔG values between 1.0 and 2.0 kcal/mol are shown in red, ΔΔG values between 0.5 and 1.0 kcal/mol are shown in light red, and ΔΔG values below 0.5 kcal/mol are shown in gray.

Loss or reduction in binding was monitored by an increase in dissociation constant K_D_ and the resulting free-energy penalty (ΔΔG) imposed by the mutation (Figure 5B). Little to no changes in ΔΔG (Figure 5, gray) were identified for residues at the edge of the basic patch, but with side chains facing away from the binding interface, namely residues K111, K136, K137, and H140. Similarly, no change in binding affinity was observed for mutants Q99A and F102A that showed a strong reduction in canonical binding in HsLARP1 and HsLARP3.[25,28,54]

Within the binding interface, a subset of mutations including K117A, H118A, N122A, K123A, and R165A had small effects on binding (Figure 5, light red), while the mutations R121A, K130A, and K139A showed a more pronounced loss in binding (Figure 5, medium red). These mutations introduced a modest free-energy penalty between 0.5 and 2.0 kcal/mol indicating a contribution of these residues to the binding affinity without the loss of a full salt bridge. This can be explained by the presence of many charged residues that can perform compensatory functions when a single mutation is introduced.

The most pronounced reduction in binding affinity occurred in mutations F114A and K166A, warranting a closer look (Figure 5, dark red). Strong NOE contacts between residues L131 and F114 to base-paired nucleotides suggest π-π stacking interaction of F114 and 5’SL RNA. Interestingly, F114 is the only residue of the canonical binding site involved in non-canonical binding. The role of K166 remains more elusive, however, the formation of a salt bridge or cation-π interaction is most likely, considering a ΔΔG over 2.0 kcal/mol.

Overall, no single-point mutation tested here entirely abolished binding. The significant involvement of charged residues however suggests a role of salt bridges in binding, most likely but not exclusively involving the stems of 5’SL that have been shown to be necessary for binding. This is evidenced through strong binding by 5’SL constructs with stem sequence mutations and our A2M5 model,[33,56] as well as stem region protection in partial nuclease digestion assays.[50]

### The La domain does not bind canonical RNA sequences or helical RNA

We tested the ability of the La domain of HsLARP6 to employ the canonical binding site for the binding of UUUUUU-3’OH (U_6_) or AAAAAA-3’OH (A_6_) that are known to bind to the canonical binding site in HsLARP3/7 and HsLARP1, respectively.[25,28,30] However, no binding was observed between the La domain for either U_6_ or A_6_ (Figure S13). This is surprising as the conserved residues determined to be necessary for binding in HsLARP1/3/7 are also present in HsLARP6 (Figure 1B), however, the La domain of HsLARP6 lacks the ability to bind these single-stranded RNAs.

To test whether the La domain binds duplex RNA indiscriminately, we used two 14-mer model hairpin RNAs and two hairpins of similar length to A2M5, named HP1 and HP2, that form a continuous helix but no bulge. Neither HP1 nor HP2 RNA bind to the La domain (Figure S14), indicating that the presence of the internal loop is essential for binding. Furthermore, the imino protons observed in intermolecular NOEs to L131 and F114 are visible in NMR spectra, indicating their participation in base-pairing. This suggests that the internal loop is structured and does not purely contribute to binding as single-stranded RNA.

## DISCUSSION

Early biochemical and biophysical studies into the binding of 5’SL RNA by the La domain of HsLARP6 suggested that only the full-length La module of HsLARP6 comprised of La domain and RRM could bind 5′SL RNA, while the La domain itself showed no 5’SL binding.[33,51] Subsequent studies by our and other laboratories however have shown that the La domain itself is capable of binding to 5’SL RNA with low nanomolar affinities.[50,55,56] The results presented here revealed how the La domain of HsLARP6 achieves high-affinity recognition of the 5′SL motif present in type I collagen mRNA by means of a non-canonical binding surface that combines electrostatic and hydrophobic interactions as well as shape complementarity. These findings substantially expand the biophysical paradigm of RNA recognition by La-related proteins, which until now was dominated by data indicating a conserved hydrophobic pocket that targets single-stranded UUU-3’OH or AAA-3’OH motifs of RNA transcripts.

### A dormant canonical binding site and emergence of a lineage-specific binding surface

Although the canonical hydrophobic binding pocket that recognizes single-stranded terminal UUU-3′OH or AAA-3′OH motifs in other LARPs is fully conserved in HsLARP6, neither U_6_ nor A_6_ oligonucleotides were found to substantially bind to the La domain (Figure S13), indicating that the canonical binding site appears to have become dormant or vestigial. From a phylogenetic standpoint, these findings indicate that the conservation of the canonical binding site co-exists with the acquisition of a lineage-specific, non-canonical binding surface that expand the La domain’s repertoire without potentially sacrificing ancestral functions. Such co-existence of dormant and specialized sites echoes observations in other families of ribonucleoproteins where evolutionary pressure favors repertoire expansion without the loss of pre-existing protein folds.[107,108]

### Biophysical determinants of 5′SL recognition

While the canonical binding in LARPs is driven mostly by π-stacking and hydrogen bonding involving a set of highly conserved residues lining the hydrophobic binding pocket (*vide supra*, Figure 1), none of these interactions were observed in the complex of 5’SL and La domain of HsLARP6. Instead, a large Lys/Arg/His-rich basic patch spanning helices α2/α3/α4 and the RNK motif provide a principal driving force via long-range Coulombic attraction to the phosphodiester backbone of the double-stranded sections of 5’SL. Mapping of the interface using CSPs, sPREs, and NOEs (Figure 4) indicated a significant overlap between the interaction site and the basic patch. Furthermore, free-energy penalties are observed when single Lys/Arg/His residues are mutated to alanine. Electrostatic patches are frequently exploited by proteins binding double-stranded RNA.[109,110] These observations underscore the electrostatic steering model of binding, in which favorable long-range Coulombic attractions stabilize the formation of the 5’SL complex.[111,112] Similar Coulombic funnels have been described for double-stranded RNA binders such as influenza A NS1 and norovirus VPg.[104,105,113]

Although the extensive basic patch appears to play a significant role in binding 5’SL RNA, its mere presence does not account for the high degree of selectivity observed in the recognition of the 5’SL motif. This is evident in binding studies showing that the La domain does not interact with continuous A-form RNA duplexes in any appreciable capacity (Figure S14). Located at the center of the electrostatic scaffold, the aromatic and aliphatic side chains of F114 and L131, respectively, form a hydrophobic patch that interacts with two adjacent, base-paired nucleotides of the internal loop as seen in NOE data (Figure 4). Furthermore, mutation of F114 to alanine incurred a free-energy penalty of over 2.0 kcal/mol, underscoring the role of the F114/L131 hydrophobic patch in binding the 5’SL RNA. Additionally, mutation of the charged residues R121A, K130A, and K139A each impose a free-energy penalty of between 1.0 and 2.0 kcal/mol, with K166A imposing a penalty of over 2.0 kcal/mol, consistent with salt bridge formation to the phosphate backbone as well as potential cation-π contacts to stacked bases. Notably, these residues surround the F114/L131 hydrophobic patch and appear to form an electrostatic “prong”, forming interactions with larger contributions than the other charged residues of the basic patch.

Binding assays with control hairpins of various lengths demonstrated that a continuous A-form duplex is insufficient for recognition and that the presence of the internal loop is essential for binding. Therefore, 5’SL recognition depends on the composite geometry of two short helices flanking a structured internal loop. The bipartite architecture of the two stems positions the internal loop and aligning it optimally with the key residues such as F114, R121, K130, and K166 while allowing the stems to engage the extended basic patch. This “3-point docking” (two stems + one internal loop) is incompatible with flexible single-stranded sequences or rigid, uninterrupted A-form helices, providing a biophysical rationale for the sharp selectivity observed *in vivo* and *in vitro*. This mechanism also reconciles the mutational tolerance of peripheral Lys/Arg residues with the more substantial nature of F114/K166 and explains how LARP6 discriminates against both poly(U/A) tails and featureless duplex RNAs.

### Concluding remarks

Overall, these results provide a complex picture of 5’SL binding with a combination of factors such as hydrophobic and electrostatic interactions paired with shape complementarity contributing to 5’SL binding. The tolerance of single-point mutations to withstand complete loss in binding provides insights into the complex binding mode but also provides a significant roadblock in describing the interaction in its entirety. This observation is further supported by the report that the La module of the invertebrate *Tetrabaena socialis* LARP6 was found to bind weakly to the human 5’SL motif that is not present in *Tetrabaena*.[114] Future high-resolution structures of the full protein-RNA complex are necessary to shed light on the specific roles of RNA nucleotides in 5’SL binding. Similarly, further studies on the full La module are required to clarify how the RRM domain contributes to this interaction and whether similar non-canonical patches are redeployed across the LARP superfamily.

## DATA AVAILABILITY

The resonance assignments of the A2M5-bound La domain of HsLARP6 have previously been published and deposited in the BMRB under accession number 31231. Atomic coordinates of A2M5-bound La domain of HsLARP6 have been deposited in the Protein Data Bank under accession code 9NGX.

## SUPPLEMENTARY DATA

Supplementary data is available at NAR online.

## AUTHOR CONTRIBUTIONS

Blaine H. Gordon: Conceptualization, Data curation, Formal analysis, Investigation, Methodology, Validation, Visualization, Writing - original draft, Writing - review and editing. Victoria S. Ogunkunle: Investigation, Writing - review and editing. Robert Silvers: Conceptualization, Data curation, Formal analysis, Funding acquisition, Methodology, Project administration, Supervision, Validation, Visualization, Writing - original draft, Writing - review and editing.

## ACKNOWLEDGEMENTS

The authors would like to thank Branko Stefanovic for insightful discussions. ARTINA had support from the European Union’s Horizon 2020 research and innovation program under the Marie Sklodowska-Curie grant agreement No. 891690. This study made use of NMRbox: National Center for Biomolecular NMR Data Processing and Analysis, a Biomedical Technology Research Resource (BTRR), which is supported by NIH grant P41GM111135 (NIGMS).

