## Supporting Information for "Non-Canonical RNA Binding of Human La-related Protein 6"

| | |
|---|---|
| Blaine H. Gordon | 0000-0002-0758-5078 |
| Victoria S. Ogunkunle | 0009-0009-0459-1331 |
| Robert Silvers | 0000-0003-0197-3878 |

### Supporting Figures

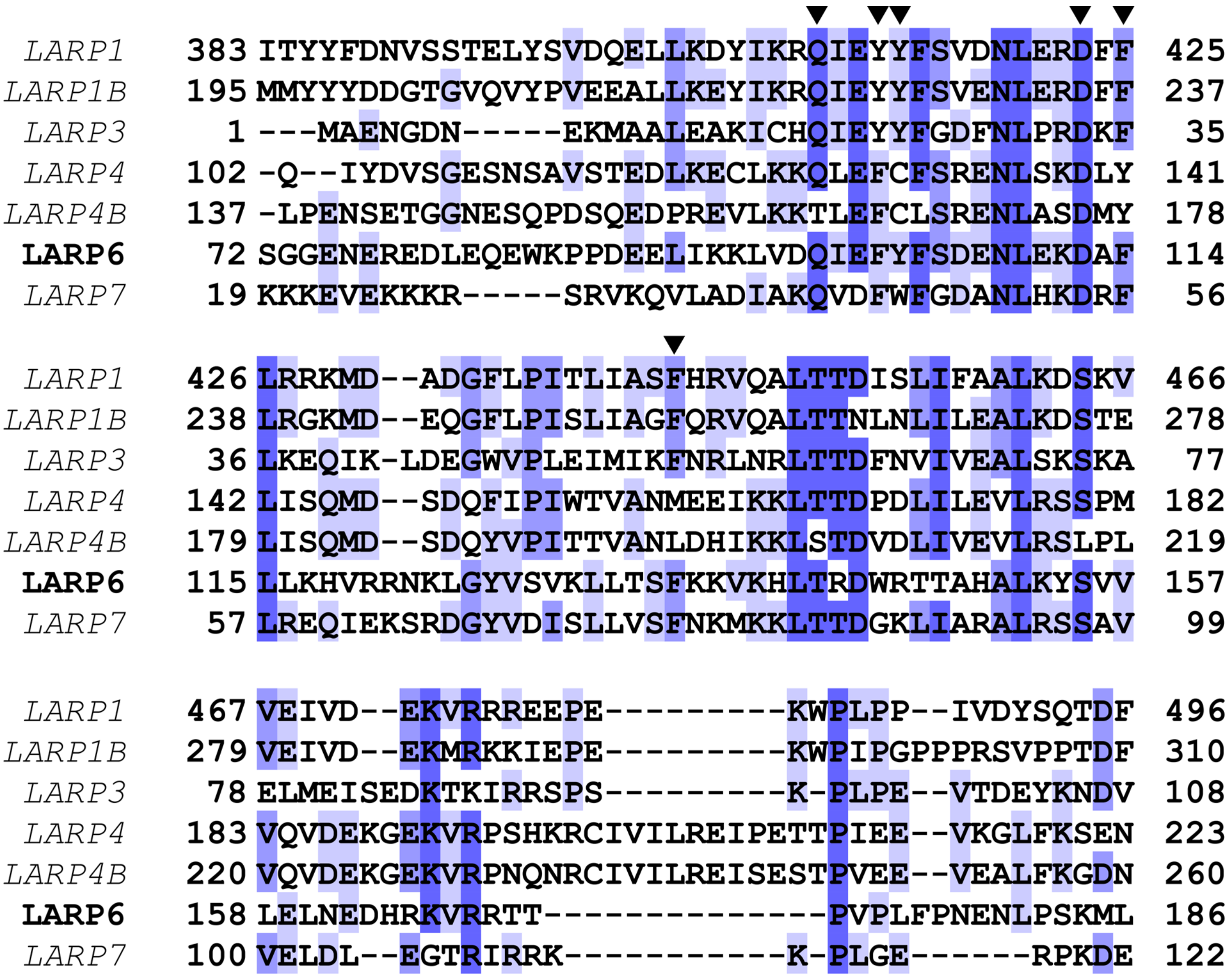


**Figure S1: Sequence alignment of the La domains of human LARPs.** Conserved residues of the canonical RNA binding pocket are indicated by inverted triangles (▼). Percentage identity is shown in shades of blue ranging from white (0%) to dark blue (100%). The following UniProt accession numbers were used: HsLARP1 (Q6PKG0-1), HsLARP1B (Q659C4-1), HsLARP3 (P05455), HsLARP4 (Q71RC2-1), HsLARP4B (Q92615), HsLARP6 (Q9BRS8-1), and HsLARP7 (Q4G0J3-1). Sequences were aligned using Clustal Omega[1] and the alignment was visualized using Jalview[2].

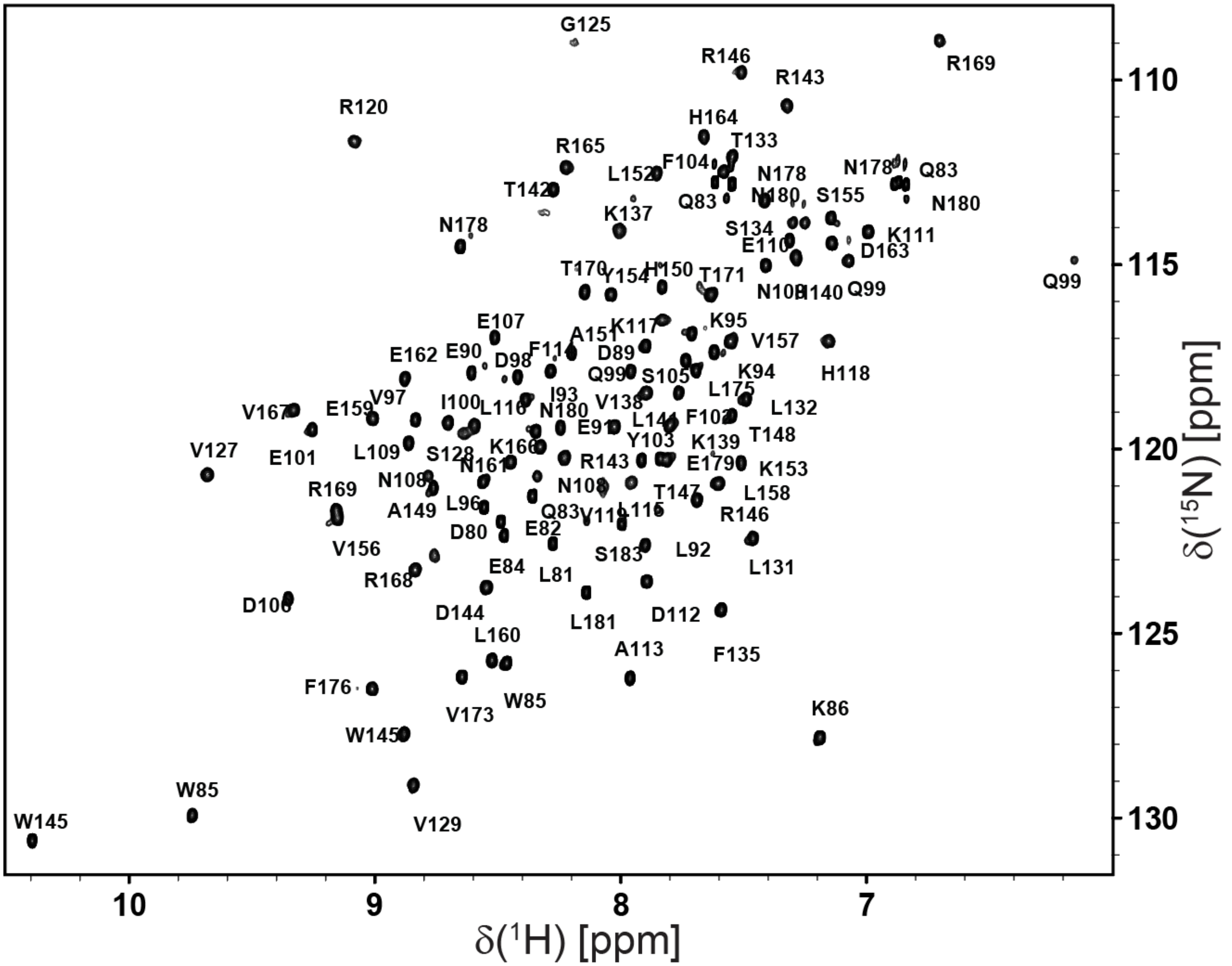


**Figure S2: Amide resonance assignment of the unbound La domain.** The [$^{1}$H, $^{15}$N]-HSQC of 500 µM [$^{15}$N]-labeled HsLARP6(79-183) was recorded at 16.4T, 298 K, and in a buffer containing 10 mM MES pH 6.5, 50 mM KCl, 10% $D_2O$, and 0.01 mg/mL DSS. The spectrum of the unbound HsLARP6(79–183) has been published previously without assignment and is reproduced here to show the full assignment.[3] Resonance assignments can be found in Table S4.

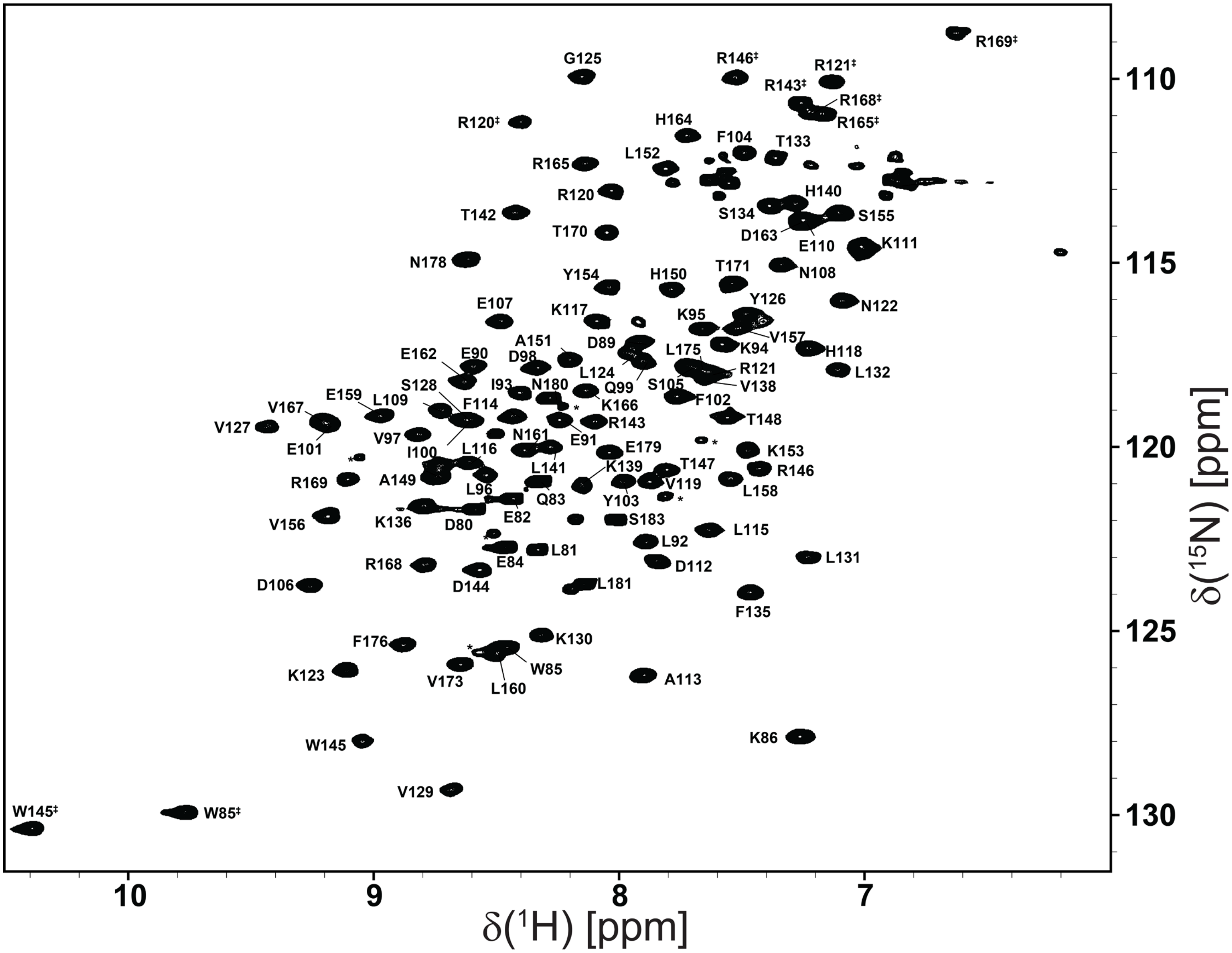


**Figure S3: Amide resonance assignment of the A2-bound La domain.** The [$^{1}$H, $^{15}$N]-HSQC of a 1:1 complex of 200 µM [$^{15}$N]-labeled HsLARP6(79-183) and natural abundance A2 RNA was recorded at 16.4T, 298 K, and in a buffer containing 10 mM MES pH 6.5, 50 mM KCl, 10% $D_2O$, and 0.01 mg/mL DSS. The spectrum of the A2-bound HsLARP6(79–183) has been published previously without assignment and is reproduced here to show the full assignment.[4] Resonance assignments can be found in Table S5.

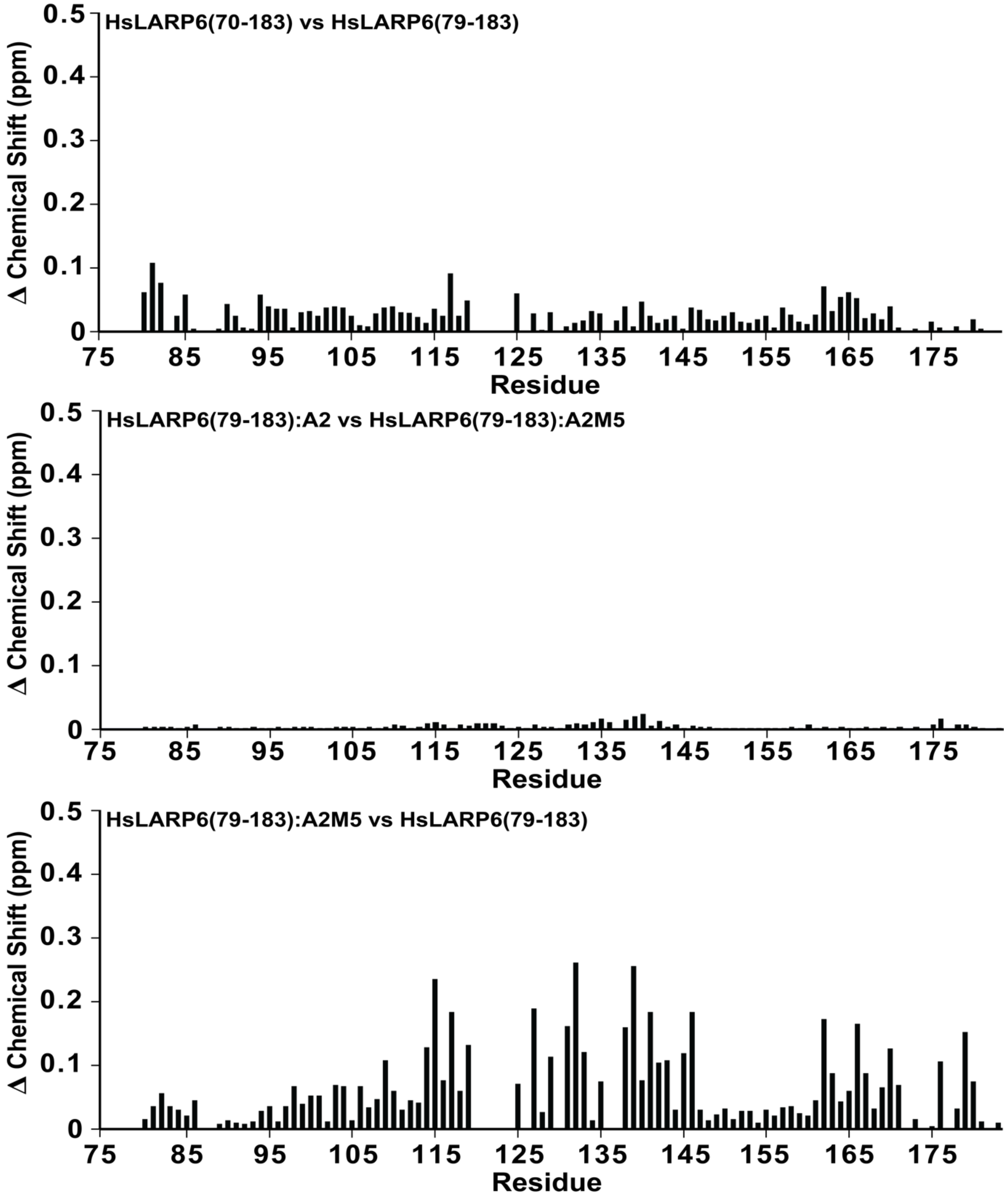


**Figure S4: Chemical shift perturbations of the La domain of HsLARP6.** Chemical shift differences were determined between backbone amide chemical shifts in [$^{15}$N]-HSQC spectra of the La domain of HsLARP6 under different conditions. CSPs between HsLARP6(79-183) and HsLARP6(70-183)[5] (top) in different NMR buffer conditions show marginal differences from alternative sequence lengths, pH and ionic strengths. HsLARP6(79-183):A2 and HsLARP6(79-183):A2M5 (middle) had the most difference in chemical shift between residues 135-140. Perturbations induced by 5'SL binding (bottom) were determined using HsLARP6(79-183) and HsLARP6(79-183):A2M5 under identical conditions. HsLARP6(79-183) and its complexes were recorded in 10 mM MES pH 6.5, 50 mM KCl, 0.01 mg/mL DSS and 10% $D_2O$.

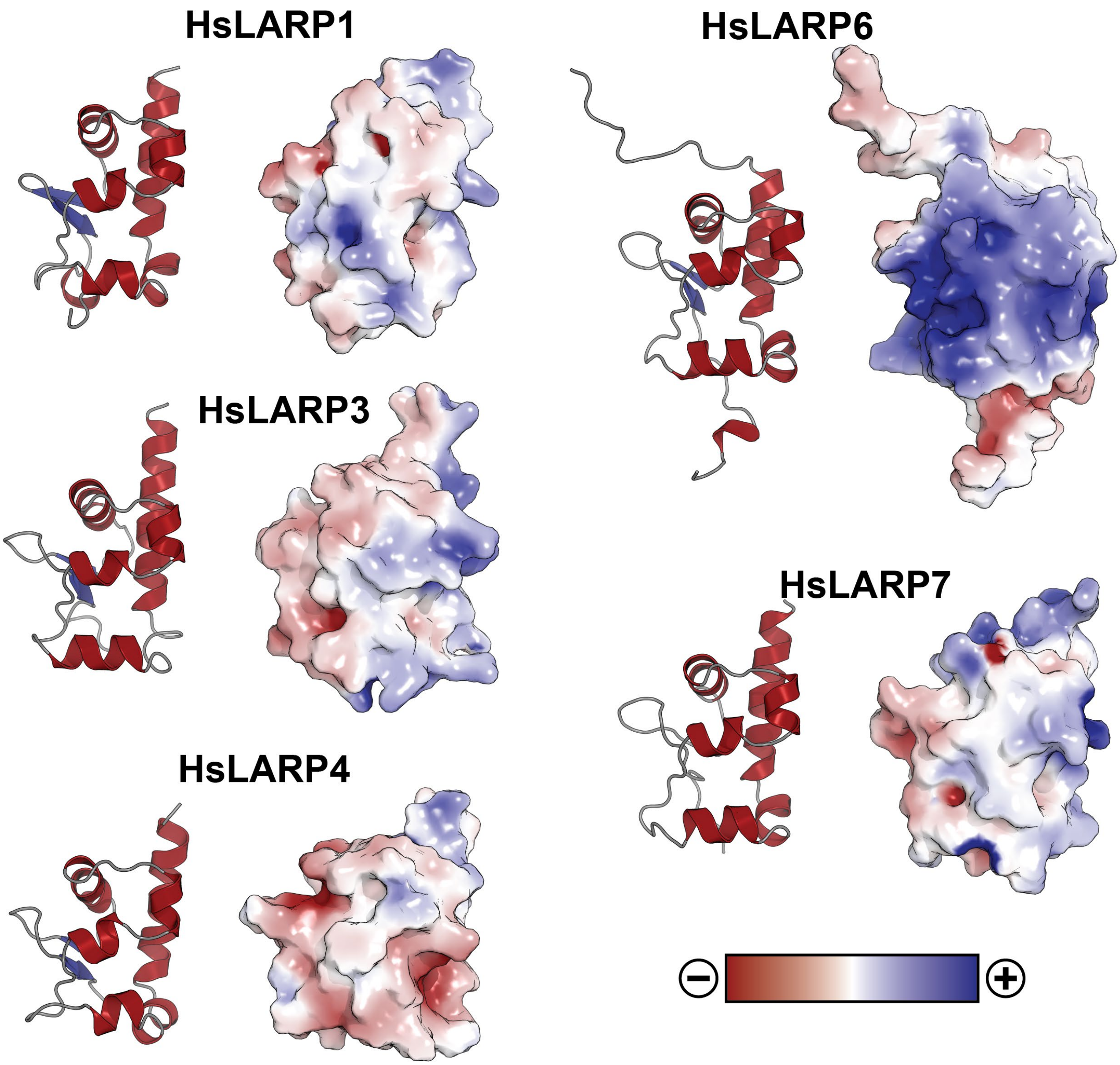


**Figure S5: Surface charges of the La domains of human LARPs.** Surface charges are shown from negative (red) to positive (blue) as determined by APBS.[6] Cartoon representations are shown for reference. The following PDB accession numbers were used: HsLARP1 (7SOR), HsLARP3 (2VOD), HsLARP4 (6I9B), HsLARP6 (9NGX), and HsLARP7 (4WKR).[7-10]

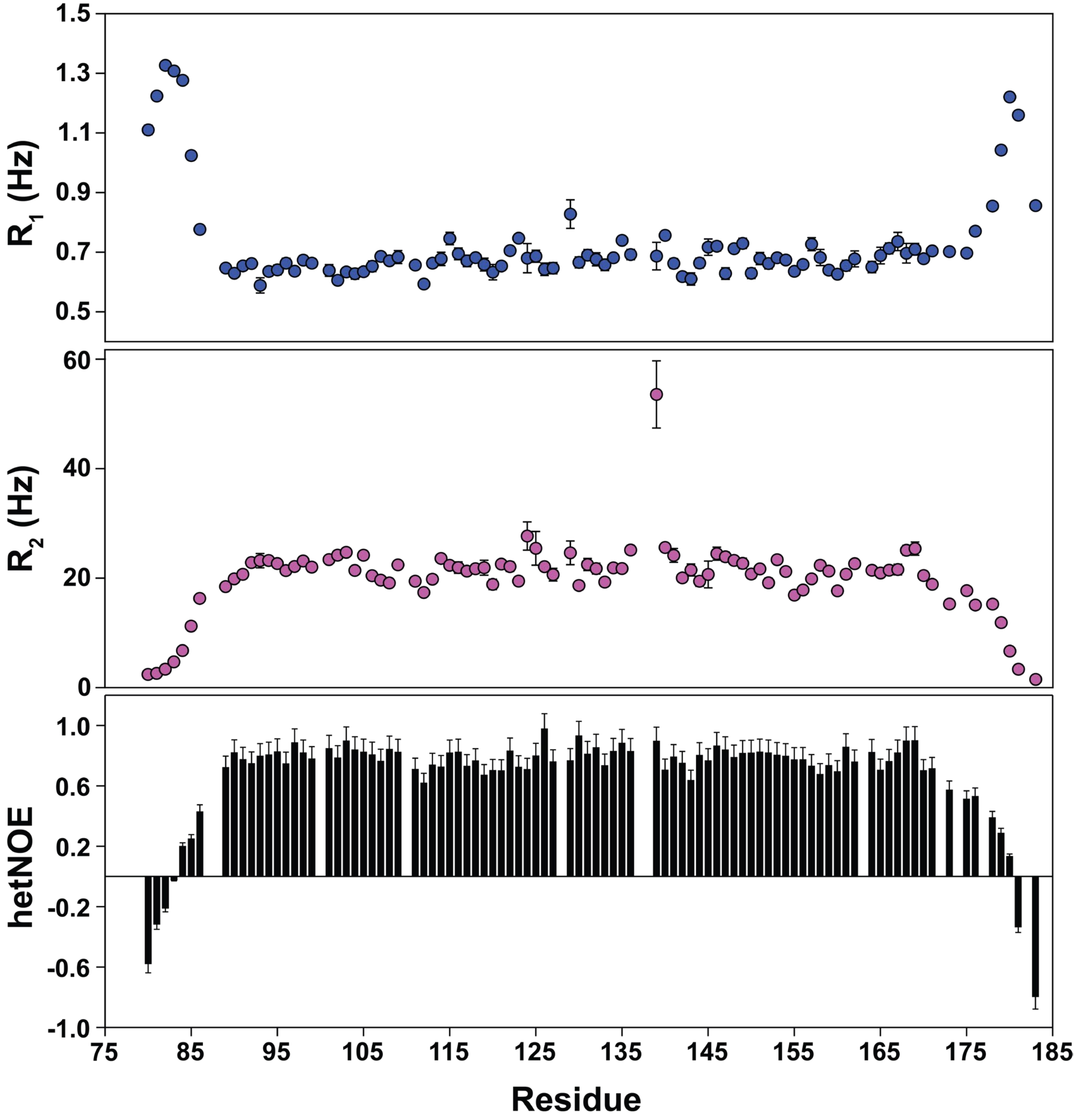


**Figure S6: Backbone amide relaxation data of the HsLARP6(79-183):A2M5 complex.** Residue-specific $^{15}$N longitudinal ($R_1$) and transverse ($R_2$) relaxation rates, as well as steady-state $^{1}$H-$^{15}$N heteronuclear nuclear Overhauser effects (hetNOE) of the HsLARP6(79-183):A2M5 complex. Errors bars reflect standard errors determined for each experiment. Relaxation rates and hetNOE data was recorded at 16.4 T and 298 K in a solution of 10 mM MES pH 6.5, 50 mM KCl, and 10% v/v $D_2O$. Relaxation data can be found in Table S7.

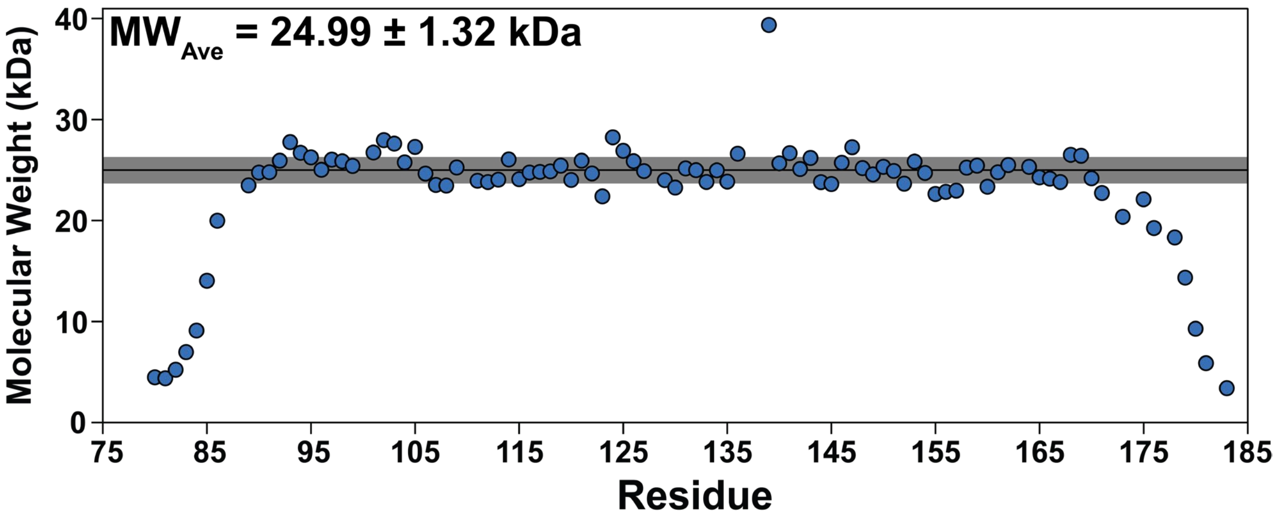


**Figure S7: Molecular weight approximation from relaxation data.** The approximate molecular weight of the HsLARP6(79-183):A2M5 complex was based on $T_1/T_2$ ratios of each residue. The average molecular weight ($MW_{Ave}$; black line) and one standard deviation (gray area) are given. Only core residues showing no large deviation in relaxation times were used in the calculation. Relaxation data can be found in Table S7.

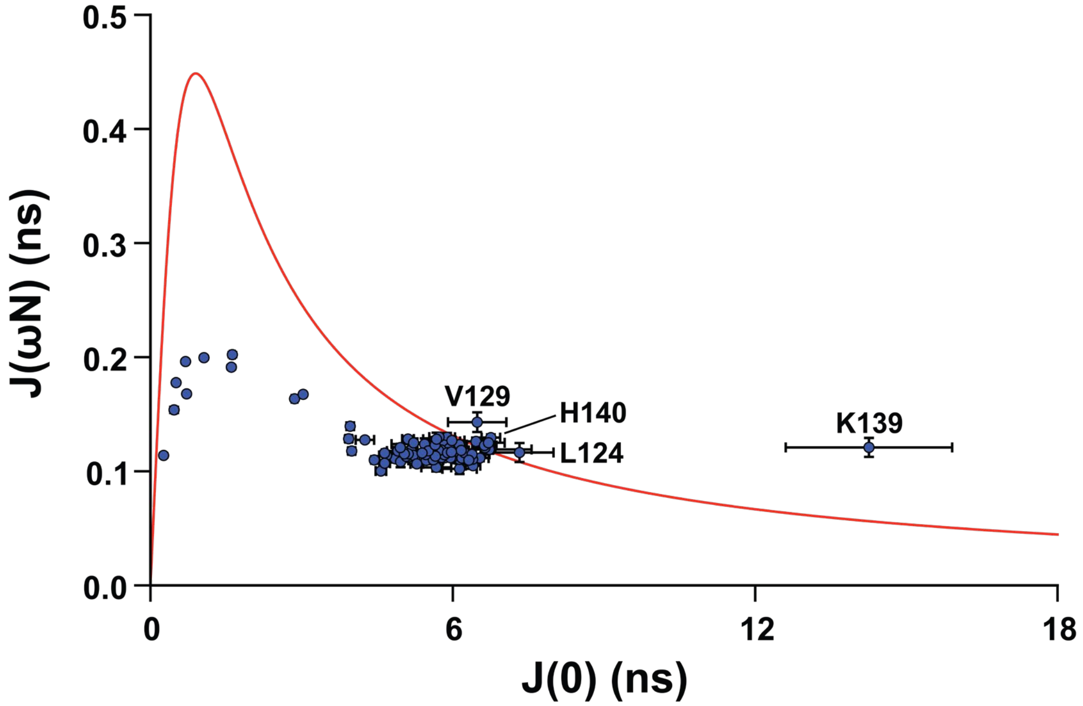


**Figure S8: Reduced spectral density function map of the HsLARP6(79-183):A2M5.** The residue-specific map of the spectral density function at J(ωN) and J(0) derived from relaxation measurements of [$^{15}$N]-HsLARP6(79-183):A2M5 is shown (blue dots). Residues extending slightly beyond or well beyond the theoretical spectral density function describing dynamics limited to isotropic global tumbling (red curve) are labeled. Error bars represent standard error propagated from the initial relaxation measurements. Relaxation data can be found in Table S7.

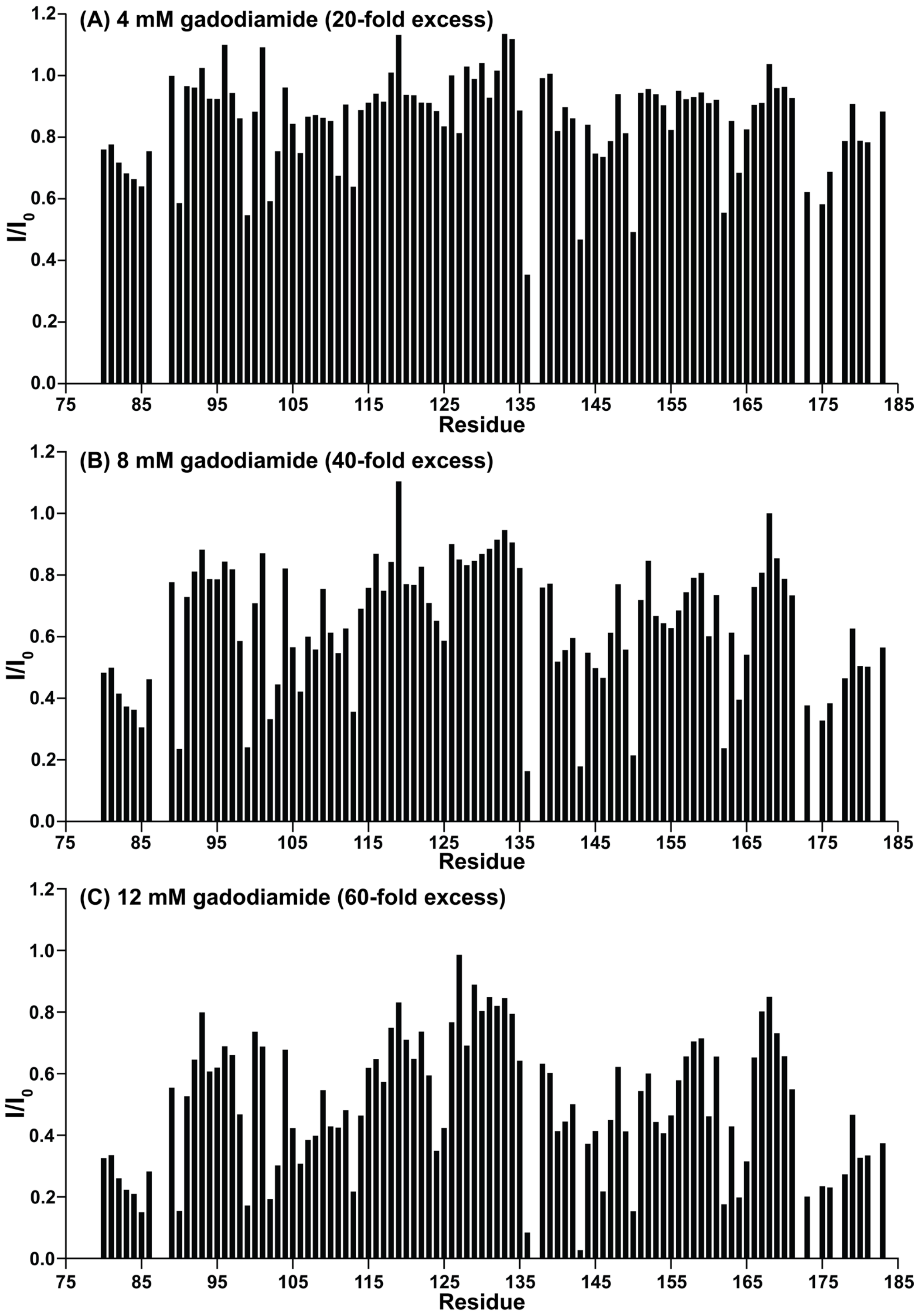


**Figure S9: Signal attenuation in the presence of paramagnetic co-solvent gadodiamide.** Signal intensities were extracted from [$^1$H, $^{15}$N]-HSQC spectra of a 1:1 complex of 200 μM [$^{15}$N]-labeled HsLARP6(79-183) and natural abundance A2 RNA recorded at 16.4T, 298 K, and in a buffer containing 10 mM MES pH 6.5, 50 mM KCl, 10% $D_2O$, and 0.01 mg/mL DSS. The intensity ratios were calculated from the spectra in the presence (I) or absence ($I_0$) of the co-solvent gadodiamide in the concentration indicated.

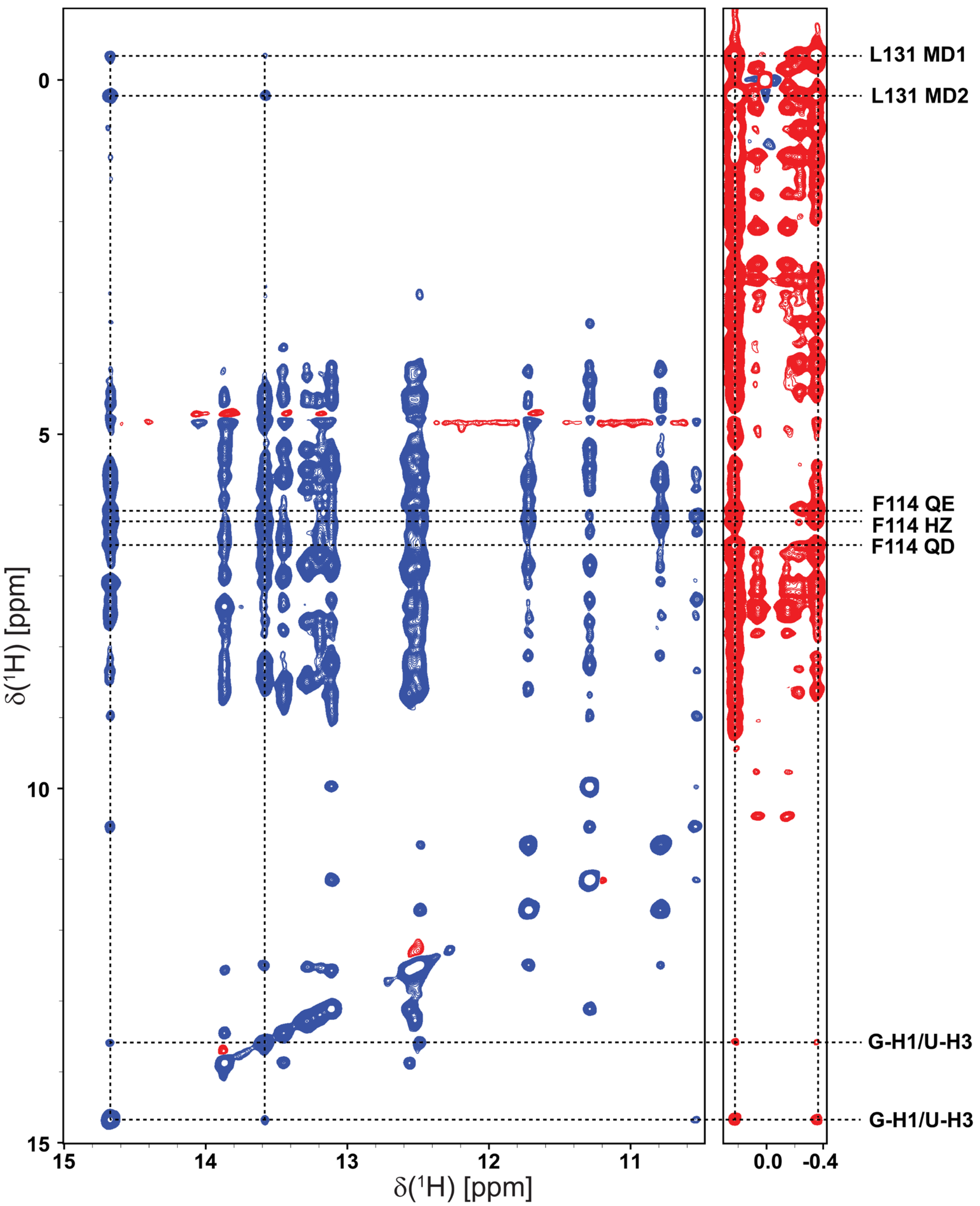


**Figure S10: Intermolecular NOE contacts between A2M5 RNA and the La domain.** The imino and methyl regions of a 2D $^1H$-$^1H$-NOESY spectrum showing intermolecular NOE contacts between A2M5 RNA and the La domain are shown. Parts of the imino region (left, blue) and methyl region (right, red) are shown. Dashed lines link diagonal and cross peaks observed in both spectra while labels indicate the identity of the observed residues from each molecule. Regions down-field and up-field of the water line have peaks with opposite signs.

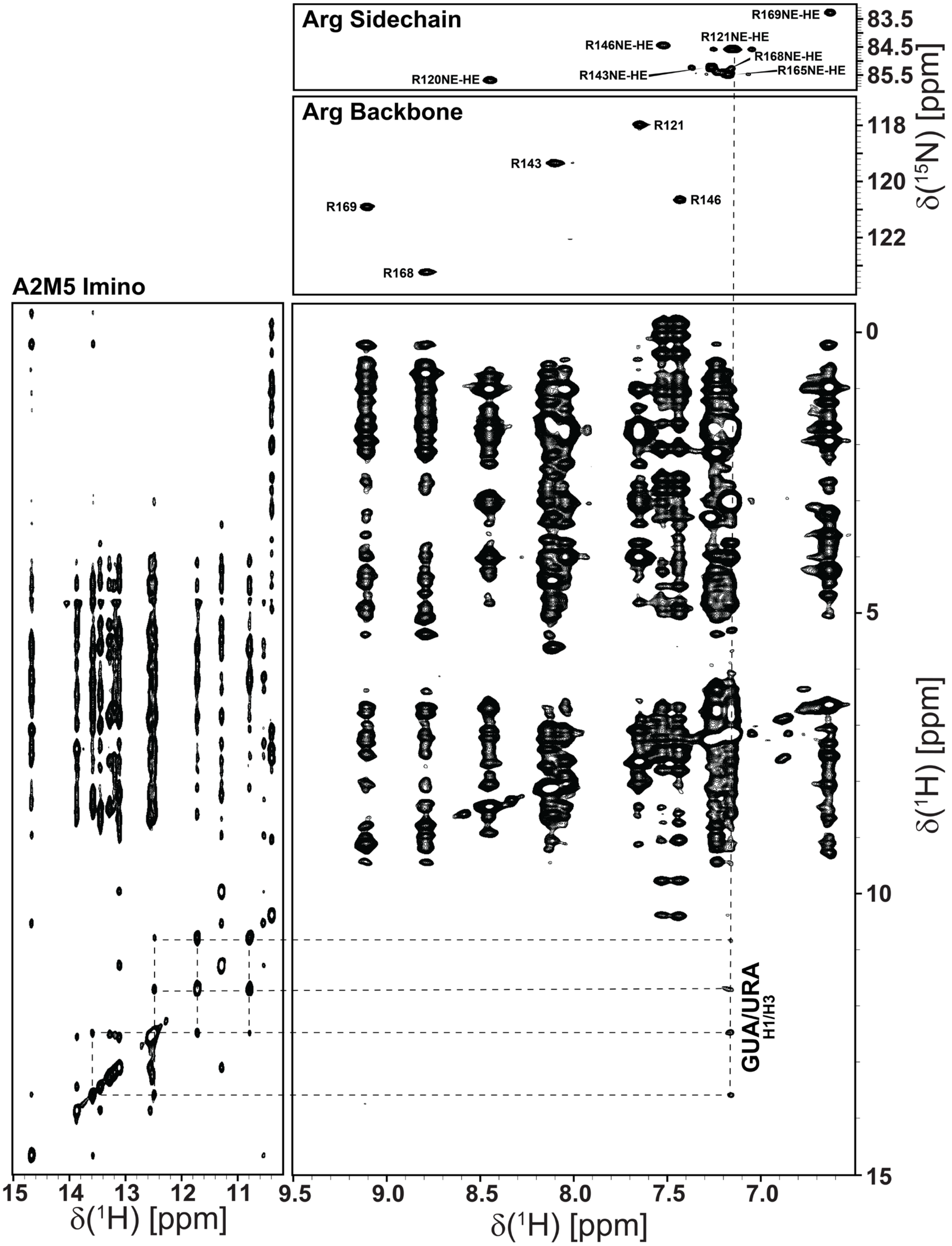


**Figure S11: Intermolecular nuclear Overhauser effects between the HsLARP6 La domain and 5'SL.** 2D $^{15}$N-edited NOESY-HSQC of [$^{13}$C,$^{15}$N-Arg]-HsLARP6(79-183) in complex with natural abundance A2M5 showing intermolecular cross peaks of arginine sidechain and RNA imino protons (bottom right). Assignments of arginine sidechain $N^E$-$H^E$ (top right) and backbone amide (middle right) resonances are given in $^{15}$N-HSQC spectra. 2D homonuclear NOESY spectrum (left) of the imino proton region of A2M5 is shown. Dashed lines connect diagonal and cross peaks present in each spectrum. All spectra were collected at 298K, 16.4 T in a buffer containing 10 mM MES pH 6.5, 50 mM KCl, 10% $D_2O$ and 0.01 mg/mL DSS. The $^{15}$N-HSQC spectra (top) have been published previously [4] and are shown here as reference.

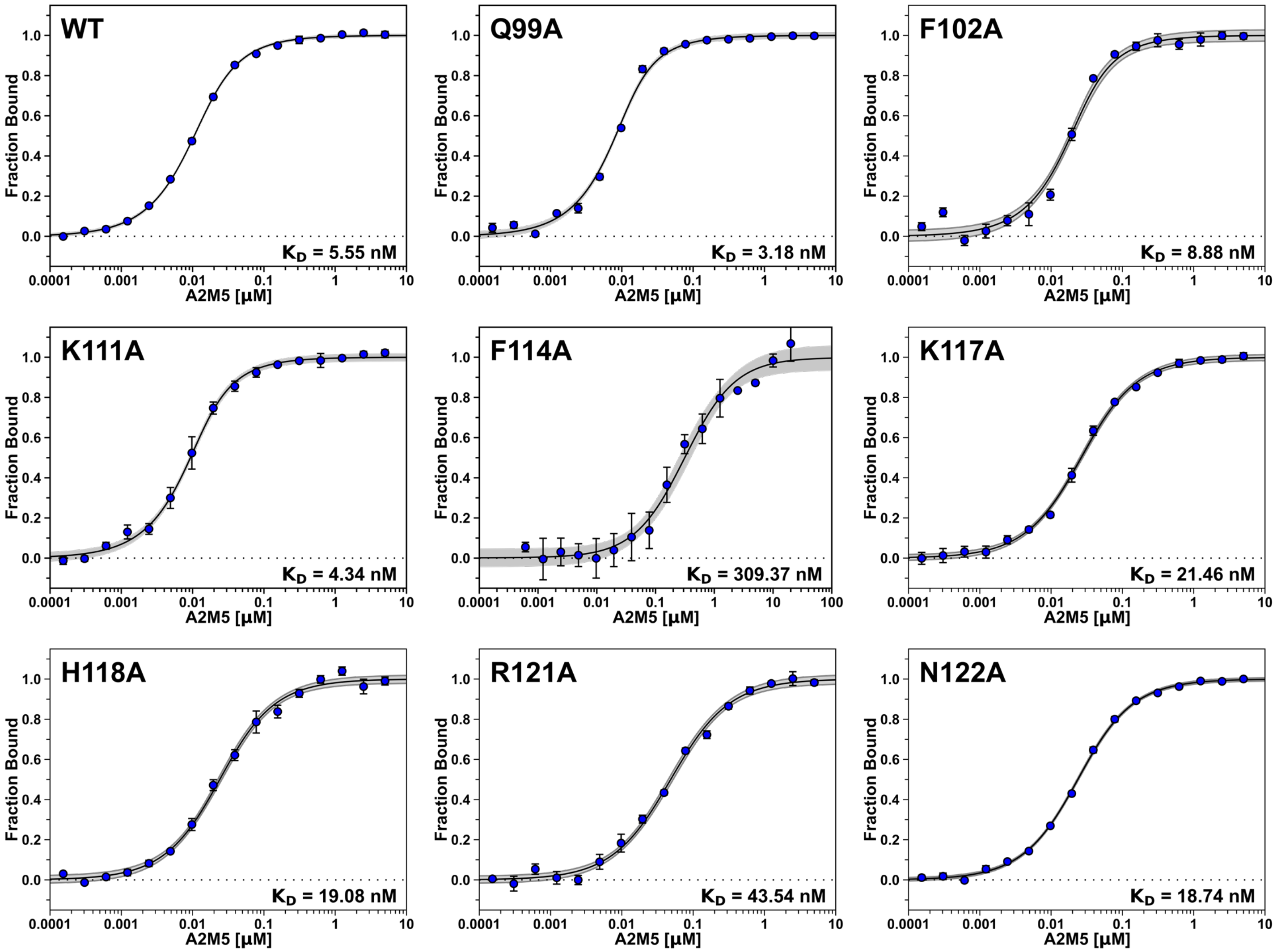


**Figure S12:** MST fitted curves of Cy5-HsLARP6 la domain and alanine point Q99A, F102A, K111A, F114A, K117A, H118A, R121A, and N122A. Dissociate constants ($K_D$s) are indicated in the bottom right of each curve. Error bars represent the standard deviation from 3 replicates of each data point. Gray areas represent the 95% confidence interval for each dataset. Data of A2M5 binding to Cy5-HsLARP6 La domain was published previously [4] and is shown here as reference.

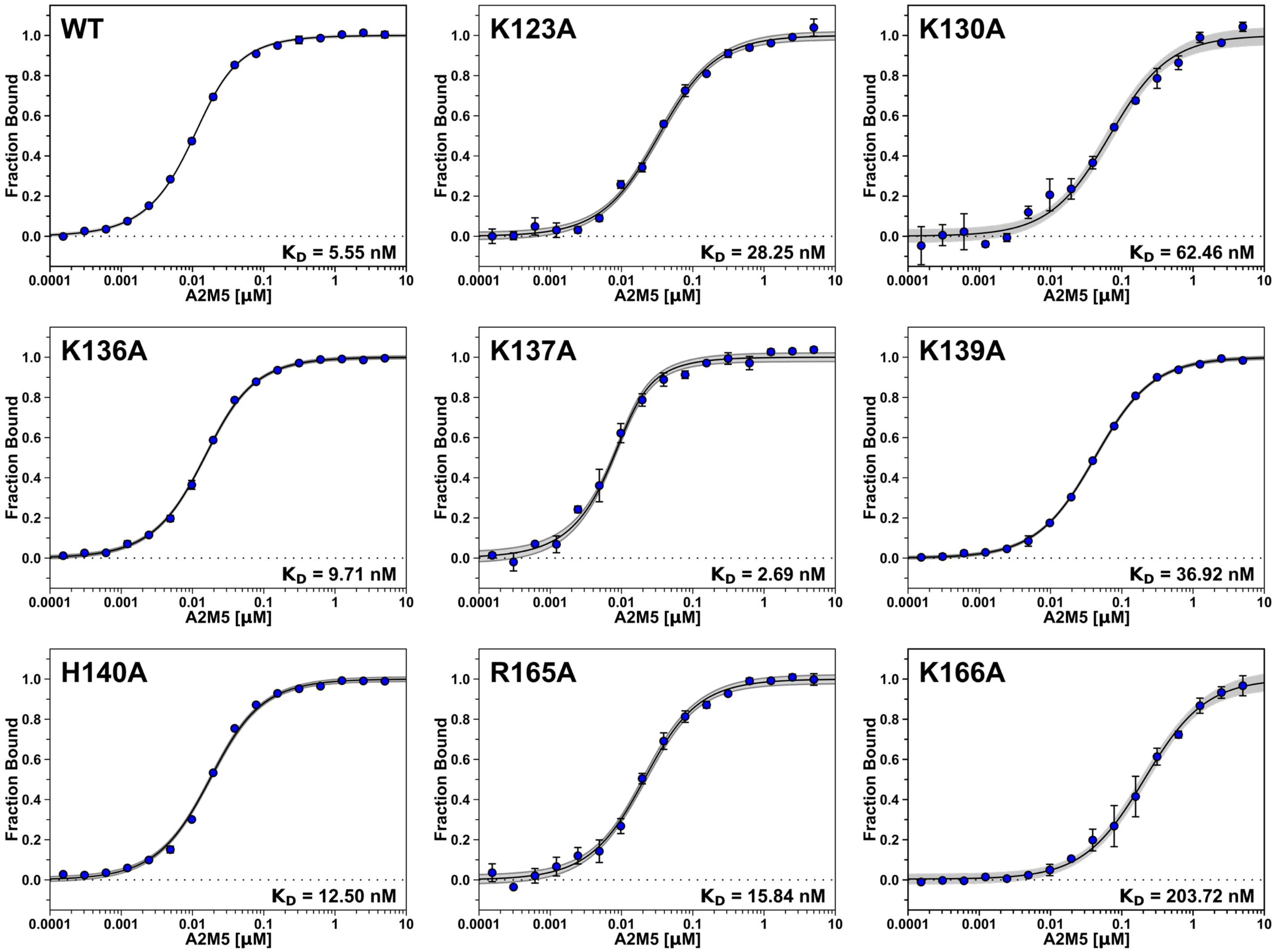


**Figure S12 (continued)** MST fitted curves of Cy5-HsLARP6 La domain and alanine points K123A, K130A, K136A, K137A, K139A, H140A, R165A, and K166A. Dissociate constants ($K_D$s) are indicated in the bottom right of each curve. Error bars represent the standard deviation from 3 replicates of each data point. Gray areas represent the 95% confidence interval for each dataset. Data of A2M5 binding to Cy5-HsLARP6 La domain was published previously [4] and is shown here as reference.

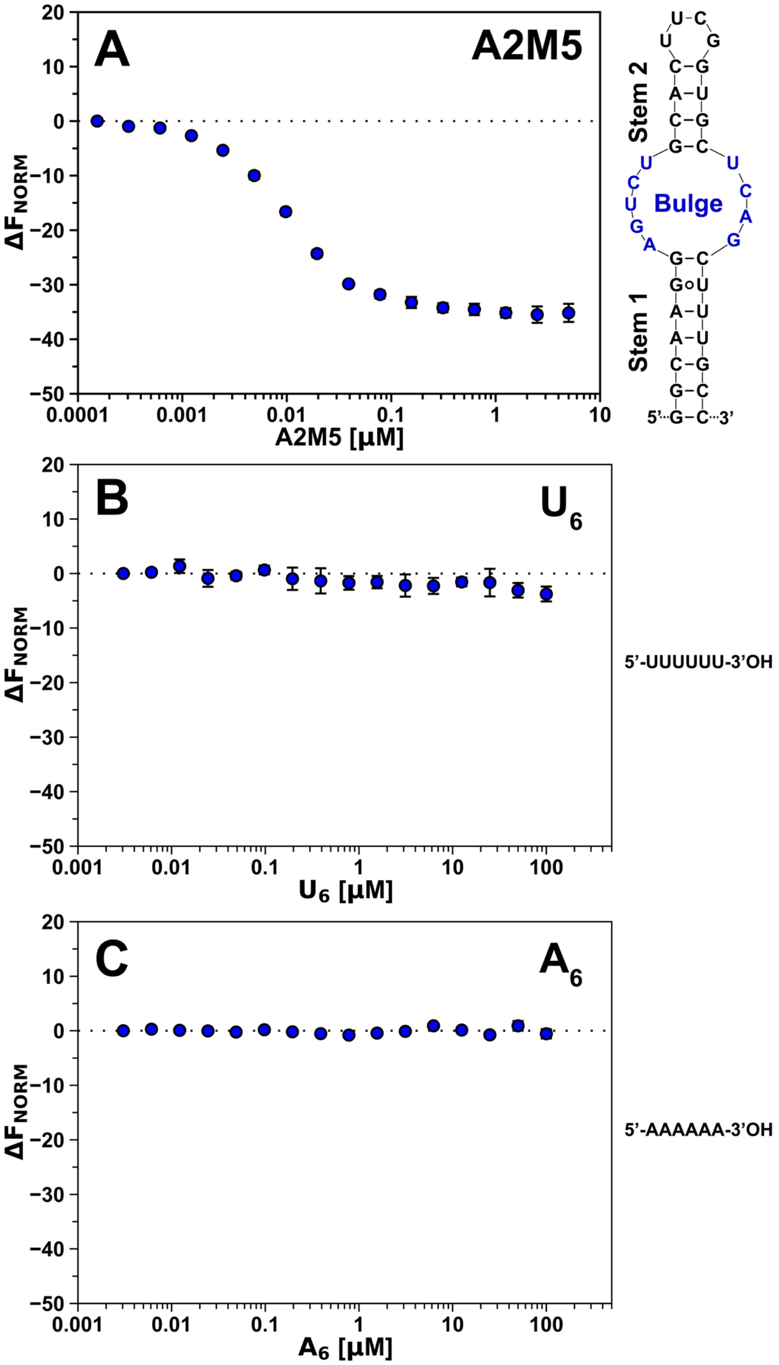


**Figure S13: Microscale thermophoresis of the HsLARP6 La domain against homopolymeric RNA:** Microscale thermophoresis (MST) of Cy5-HsLARP6 in the presence of (A) A2M5, (B) poly(U), and (C) poly(A). RNA sequences and predicted secondary structures are presented on the right of each panel. Error bars represent the standard deviation from 3 replicates of each data point. Data of A2M5 binding to Cy5-HsLARP6 La domain was published previously [4] and is shown here as reference.

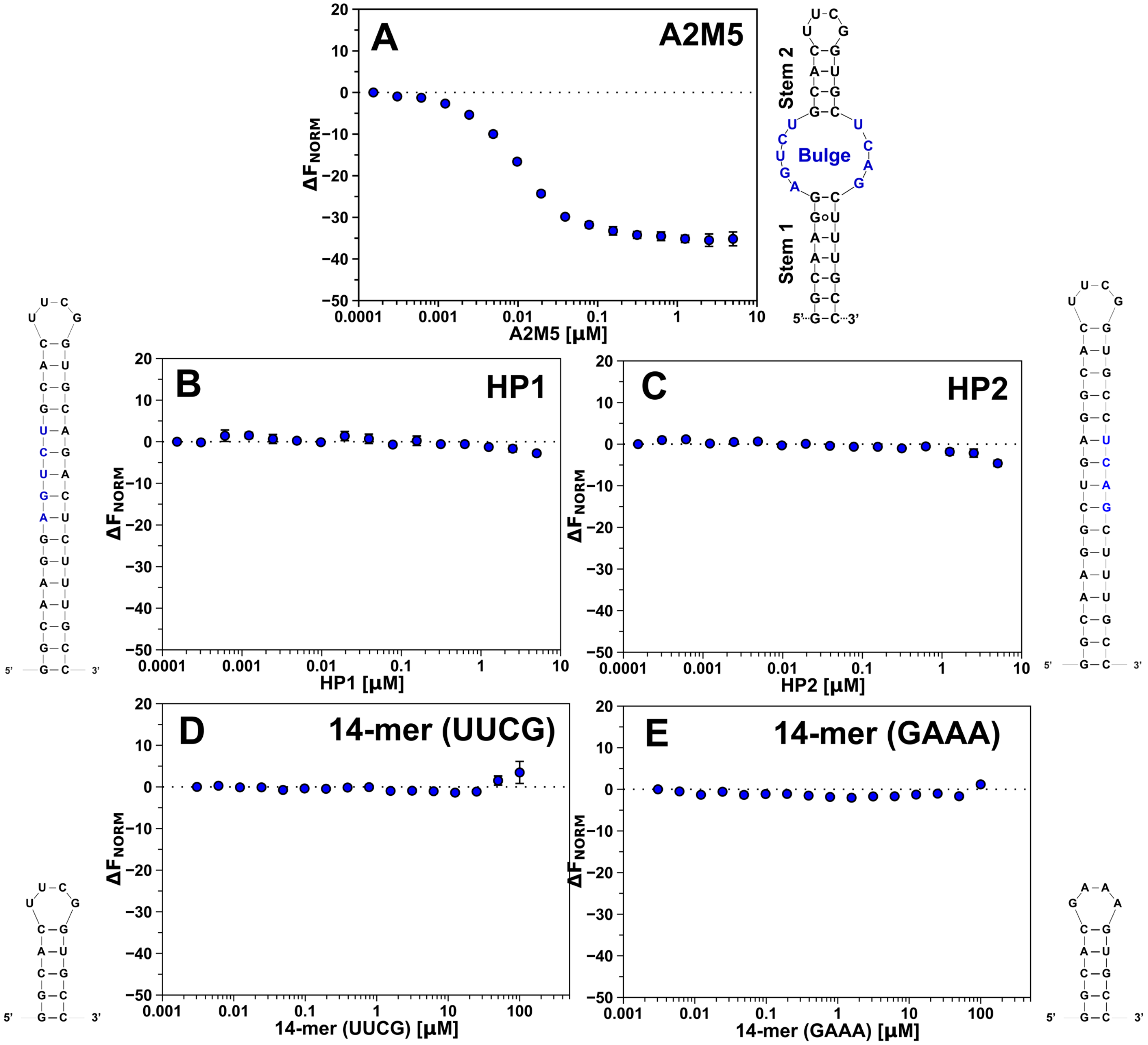


**Figure S14: Microscale thermophoresis of the HsLARP6 La domain against hairpin RNA:** MST of Cy5-HsLARP6 La domain in the presence of (A) A2M5, (B) hairpin 1 (HP1), (C) hairpin 2 (HP2), (D) UUCG tetraloop 14-mer, and (E) GAAA tetraloop 14-mer. RNA sequences and predicted secondary structures are presented adjacent of each panel. Residues from the sequence of the internal loop are show in blue. Data of A2M5 binding to Cy5-HsLARP6 La domain was published previously [4] and is shown here as reference.

### Supporting Tables

**Table S1:** Sequences of proteins

| Construct Name | Sequence |
|---|---|
| **His$_6$-TEV-HsLARP6(79-183)** | MGHHHHHHGSGSENLYFQGEDLEQEWKPPDEELIKKLVDQIEFYFSDENLEKDAFLLKHVRRNKLGYVSVKLLT<br>SFKKVKHLTRDWRTTAHALKYSVVLELNEDHRKVRRTTPVPLFPNENLPS |
| **HsLARP6(79-183)** | GEDLEQEWKPPDEELIKKLVDQIEFYFSDENLEKDAFLLKHVRRNKLGYVSVKLLTSFKKVKHLTRDWRTTAHA<br>LKYSVVLELNEDHRKVRRTTPVPLFPNENLPS |
| **His$_6$-TEV-HsLARP6(73-183)G74C** | MGHHHHHHGSGSENLYFQGCENEREDLEQEWKPPDEELIKKLVDQIEFYFSDENLEKDAFLLKHVRRNKLGYVS<br>VKLLTSFKKVKHLTRDWRTTAHALKYSVVLELNEDHRKVRRTTPVPLFPNENLPS |
| **Cy5-HsLAPR6(73-183)[a,b]** | MGHHHHHHGSGSENLYFQG**C**ENEREDLEQEWKPPDEELIKKLVDQIEFYFSDENLEKDAFLLKHVRRNKLGYVS<br>VKLLTSFKKVKHLTRDWRTTAHALKYSVVLELNEDHRKVRRTTPVPLFPNENLPS |
| **Cy5-HsLAPR6(73-183)Q99A[a,c]** | MGHHHHHHGSGSENLYFQG**C**ENEREDLEQEWKPPDEELIKKLVD**A**IEFYFSDENLEKDAFLLKHVRRNKLGYVS<br>VKLLTSFKKVKHLTRDWRTTAHALKYSVVLELNEDHRKVRRTTPVPLFPNENLPS |
| **Cy5-HsLAPR6(73-183)F102A[a,c]** | MGHHHHHHGSGSENLYFQG**C**ENEREDLEQEWKPPDEELIKKLVDQIE**A**YFSDENLEKDAFLLKHVRRNKLGYVS<br>VKLLTSFKKVKHLTRDWRTTAHALKYSVVLELNEDHRKVRRTTPVPLFPNENLPS |
| **Cy5-HsLAPR6(73-183)K111A[a,c]** | MGHHHHHHGSGSENLYFQG**C**ENEREDLEQEWKPPDEELIKKLVDQIEFYFSDENLE**A**DAFLLKHVRRNKLGYVS<br>VKLLTSFKKVKHLTRDWRTTAHALKYSVVLELNEDHRKVRRTTPVPLFPNENLPS |
| **Cy5-HsLAPR6(73-183)F114A[a,c]** | MGHHHHHHGSGSENLYFQG**C**ENEREDLEQEWKPPDEELIKKLVDQIEFYFSDENLEKDA**A**LLKHVRRNKLGYVS<br>VKLLTSFKKVKHLTRDWRTTAHALKYSVVLELNEDHRKVRRTTPVPLFPNENLPS |
| **Cy5-HsLAPR6(73-183)K117A[a,c]** | MGHHHHHHGSGSENLYFQG**C**ENEREDLEQEWKPPDEELIKKLVDQIEFYFSDENLEKDAFLL**A**HVRRNKLGYVS<br>VKLLTSFKKVKHLTRDWRTTAHALKYSVVLELNEDHRKVRRTTPVPLFPNENLPS |
| **Cy5-HsLAPR6(73-183)H118A[a,c]** | MGHHHHHHGSGSENLYFQG**C**ENEREDLEQEWKPPDEELIKKLVDQIEFYFSDENLEKDAFLLK**A**VRRNKLGYVS<br>VKLLTSFKKVKHLTRDWRTTAHALKYSVVLELNEDHRKVRRTTPVPLFPNENLPS |
| **Cy5-HsLAPR6(73-183)R121A[a,c]** | MGHHHHHHGSGSENLYFQG**C**ENEREDLEQEWKPPDEELIKKLVDQIEFYFSDENLEKDAFLLKHVR**A**NKLGYVS<br>VKLLTSFKKVKHLTRDWRTTAHALKYSVVLELNEDHRKVRRTTPVPLFPNENLPS |
| **Cy5-HsLAPR6(73-183)N122A[a,c]** | MGHHHHHHGSGSENLYFQG**C**ENEREDLEQEWKPPDEELIKKLVDQIEFYFSDENLEKDAFLLKHVRR**A**KLGYVS<br>VKLLTSFKKVKHLTRDWRTTAHALKYSVVLELNEDHRKVRRTTPVPLFPNENLPS |
| **Cy5-HsLAPR6(73-183)K123A[a,c]** | MGHHHHHHGSGSENLYFQG**C**ENEREDLEQEWKPPDEELIKKLVDQIEFYFSDENLEKDAFLLKHVRRN**A**LGYVS<br>VKLLTSFKKVKHLTRDWRTTAHALKYSVVLELNEDHRKVRRTTPVPLFPNENLPS |
| **Cy5-HsLAPR6(73-183)K130A[a,c]** | MGHHHHHHGSGSENLYFQG**C**ENEREDLEQEWKPPDEELIKKLVDQIEFYFSDENLEKDAFLLKHVRRNKLGYVS<br>V**A**LLTSFKKVKHLTRDWRTTAHALKYSVVLELNEDHRKVRRTTPVPLFPNENLPS |
| **Cy5-HsLAPR6(73-183)K136A[a,c]** | MGHHHHHHGSGSENLYFQG**C**ENEREDLEQEWKPPDEELIKKLVDQIEFYFSDENLEKDAFLLKHVRRNKLGYVS<br>VKLLTSF**A**KVKHLTRDWRTTAHALKYSVVLELNEDHRKVRRTTPVPLFPNENLPS |
| **Cy5-HsLAPR6(73-183)K137A[a,c]** | MGHHHHHHGSGSENLYFQG**C**ENEREDLEQEWKPPDEELIKKLVDQIEFYFSDENLEKDAFLLKHVRRNKLGYVS<br>VKLLTSFK**A**VKHLTRDWRTTAHALKYSVVLELNEDHRKVRRTTPVPLFPNENLPS |
| **Cy5-HsLAPR6(73-183)K139A[a,c]** | MGHHHHHHGSGSENLYFQG**C**ENEREDLEQEWKPPDEELIKKLVDQIEFYFSDENLEKDAFLLKHVRRNKLGYVS<br>VKLLTSFKKV**A**HLTRDWRTTAHALKYSVVLELNEDHRKVRRTTPVPLFPNENLPS |
| **Cy5-HsLAPR6(73-183)H140A[a,c]** | MGHHHHHHGSGSENLYFQG**C**ENEREDLEQEWKPPDEELIKKLVDQIEFYFSDENLEKDAFLLKHVRRNKLGYVS<br>VKLLTSFKKVK**A**LTRDWRTTAHALKYSVVLELNEDHRKVRRTTPVPLFPNENLPS |
| **Cy5-HsLAPR6(73-183)R165A[a,c]** | MGHHHHHHGSGSENLYFQG**C**ENEREDLEQEWKPPDEELIKKLVDQIEFYFSDENLEKDAFLLKHVRRNKLGYVS<br>VKLLTSFKKVKHLTRDWRTTAHALKYSVVLELNEDH**A**KVRRTTPVPLFPNENLPS |
| **Cy5-HsLAPR6(73-183)K166A[a,c]** | MGHHHHHHGSGSENLYFQG**C**ENEREDLEQEWKPPDEELIKKLVDQIEFYFSDENLEKDAFLLKHVRRNKLGYVS<br>VKLLTSFKKVKHLTRDWRTTAHALKYSVVLELNEDHR**A**VRRTTPVPLFPNENLPS |

[a] The Cy5-labeled cysteine is shown in bold

[b] This Cy5-labeled protein is referred to as wild type despite its G74C mutation in the N-terminus. All residues in the core domain are unaltered and carry no mutations.

[c] This Cy5-labeled protein carries a single alanine mutation in the core domain. The position is shown in bold and is colored red.

**Table S2:** Sequences of DNA oligonucleotides

| Construct Name | Sequence |
|---|---|
| **T7 Promoter** | 5'-TAATACGACTCACTATAG-3' |
| **A2 Template[a]** | 5'-mCmACAAAGCTGAGCATGTCTAGCACTTAGACATGCAGACTCCTTGTGCCTATAGTGAGTCGTATTACAT-3' |
| **A2M5 Template[a]** | 5'-mGmGCAAAGCTGAGCACCGAAGTGCAGACTCCTTGCCTATAGTGAGTCGTATTA-3' |
| **HP1 Template[a]** | 5'-mGmGCAAAGAGTCTGCACCGAAGTGCAGACTCCTTGCCTATAGTGAGTCGTATTA-3' |
| **HP2 Template[a]** | 5'-mGmGCAAAGCTGAGGCACCGAAGTGCCTCAGCCTTGCCTATAGTGAGTCGTATTA-3' |

[a] Template DNA carried 2'O-methylated nucleotides in position 1 and 2 to reduce untemplated nucleotide additions by T7 RNA polymerase during *in vitro* transcription.

**Table S3:** Sequences of RNA oligonucleotides

| Construct Name | Sequence |
|---|---|
| **A2[a,c]** | 5'-GGCACAAGG**AGUCU**GCAUGUCUAAGUGCUAGACAUGC**UCAG**CUUUGUG-3' |
| **A2M5[a,c]** | 5'-GGCAAGG**AGUCU**GCACUUCGGUGC**UCAG**CUUUGCC-3' |
| **HP1[a,c]** | 5'-GGCAAGG**AGUCU**GCACUUCGGUGCAGACUCUUUGCC-3' |
| **HP2[a,c]** | 5'-GGCAAGGCUGAGGCACUUCGGUGCC**UCAG**CUUUGCC-3' |
| **14-mer (UUCG)[b,c]** | 5'-GGCACUUCGGUGCC-3' |
| **14-mer (GAAA)[b,c]** | 5'-GGCACGAAAGUGCC-3' |
| **$A_6$[b]** | 5'-AAAAAA-3' |
| **$U_6$[b]** | 5'-UUUUUU-3' |

[a] RNA was produced by *in vitro* transcription from DNA templates (see Table S2)

[b] RNA was produced by phosphoramidite synthesis and ordered as RNase-free and HPLC-purified oligonucleotides from GenScript Corp., USA

[c] The position of internal loop or hairpin loop nucleotides are shown in bold or underlined, respectively.

**Table S4:** Chemical shifts of unbound HsLARP6(79-183) at 298 K in buffer containing 10 mM MES pH 6.5, 50 mM KCl, 10% $D_2O$, and 0.01 mg/mL DSS

| Residue | δ($^1$H) [ppm] | δ($^{15}$N) [ppm] |
|---|---|---|
| D80 | 8.556 | 121.576 |
| L81 | 8.276 | 122.559 |
| E82 | 8.487 | 121.968 |
| Q83 | 8.359 | 121.278 |
| E84 | 8.474 | 122.339 |
| W85 | 8.464 | 125.799 |
| K86 | 7.189 | 127.816 |
| P87 | - | - |
| P88 | - | - |
| D89 | 7.897 | 117.201 |
| E90 | 8.606 | 117.932 |
| E91 | 8.244 | 119.415 |
| L92 | 7.900 | 122.604 |
| I93 | 8.386 | 118.664 |
| K94 | 7.619 | 117.363 |
| K95 | 7.710 | 116.871 |
| L96 | 8.559 | 120.886 |
| V97 | 8.836 | 119.205 |
| D98 | 8.419 | 118.044 |
| Q99 | 7.959 | 117.892 |
| I100 | 8.702 | 119.286 |
| E101 | 9.255 | 119.474 |
| F102 | 7.764 | 118.470 |
| Y103 | 7.915 | 120.298 |
| F104 | 7.581 | 112.492 |
| S105 | 7.735 | 117.600 |
| D106 | 9.353 | 124.053 |
| E107 | 8.511 | 116.973 |
| N108 | 7.409 | 115.022 |
| L109 | 8.862 | 119.842 |
| E110 | 7.313 | 114.348 |
| K111 | 6.991 | 114.107 |
| D112 | 7.893 | 123.586 |
| A113 | 7.962 | 126.207 |
| F114 | 8.285 | 117.888 |
| L115 | 7.956 | 120.902 |
| L116 | 8.595 | 119.363 |
| K117 | 7.830 | 116.499 |
| H118 | 7.154 | 117.072 |
| V119 | 8.072 | 121.055 |
| R120 | - | - |
| R121 | - | - |
| N122 | - | - |
| K123 | - | - |
| L124 | - | - |
| G125 | 8.185 | 108.970 |
| Y126 | - | - |
| V127 | 9.680 | 120.698 |
| S128 | 8.633 | 119.574 |
| V129 | 8.843 | 129.107 |
| K130 | - | - |
| L131 | 7.461 | 122.423 |

| Residue | δ($^1$H) [ppm] | δ($^{15}$N) [ppm] |
|---|---|---|
| L132 | 7.489 | 118.647 |
| T133 | 7.545 | 112.077 |
| S134 | 7.414 | 113.262 |
| F135 | 7.589 | 124.356 |
| K136 | - | - |
| K137 | 8.006 | 114.075 |
| V138 | 7.896 | 118.479 |
| K139 | 7.799 | 119.358 |
| H140 | 7.286 | 114.808 |
| L141 | 8.025 | 119.395 |
| T142 | 8.273 | 112.961 |
| R143 | 8.228 | 120.235 |
| D144 | 8.546 | 123.752 |
| W145 | 8.880 | 127.720 |
| R146 | 7.689 | 121.379 |
| T147 | 7.838 | 120.263 |
| T148 | 7.546 | 119.089 |
| A149 | 8.762 | 121.054 |
| H150 | 7.830 | 115.608 |
| A151 | 8.200 | 117.397 |
| L152 | 7.852 | 112.524 |
| K153 | 7.509 | 120.374 |
| Y154 | 8.040 | 115.813 |
| S155 | 7.144 | 113.735 |
| V156 | 9.153 | 121.828 |
| V157 | 7.553 | 117.082 |
| L158 | 7.603 | 120.934 |
| E159 | - | - |
| L160 | 8.524 | 125.727 |
| N161 | 8.447 | 120.349 |
| E162 | 8.878 | 118.090 |
| D163 | 7.141 | 114.421 |
| H164 | 7.661 | 111.536 |
| R165 | 8.223 | 112.364 |
| K166 | 8.327 | 119.940 |
| V167 | 9.329 | 118.943 |
| R168 | 8.836 | 123.275 |
| R169 | 9.156 | 121.712 |
| T170 | 8.146 | 115.746 |
| T171 | 7.632 | 115.817 |
| P172 | - | - |
| V173 | 8.645 | 126.173 |
| P174 | - | - |
| L175 | 7.693 | 117.870 |
| F176 | 9.011 | 126.498 |
| P177 | - | - |
| N178 | 8.652 | 114.508 |
| E179 | 7.811 | 120.290 |
| N180 | 8.344 | 119.517 |
| L181 | 8.140 | 123.889 |
| P182 | - | - |
| S183 | 7.996 | 122.027 |

**Table S5:** Chemical shifts of the 1:1 complex of HsLARP6(79-183) and A2 RNA in at 298 K in buffer containing 10 mM MES pH 6.5, 50 mM KCl, 10% $D_2O$, and 0.01 mg/mL DSS

| Residue | δ($^1$H) [ppm] | δ($^{15}$N) [ppm] |
|---|---|---|
| D80 | 8.582 | 121.691 |
| L81 | 8.330 | 122.791 |
| E82 | 8.430 | 121.407 |
| Q83 | 8.319 | 120.936 |
| E84 | 8.459 | 122.723 |
| W85 | 8.455 | 125.450 |
| K86 | 7.263 | 127.867 |
| P87 | - | - |
| P88 | - | - |
| D89 | 7.916 | 117.150 |
| E90 | 8.594 | 117.809 |
| E91 | 8.242 | 119.271 |
| L92 | 7.889 | 122.573 |
| I93 | 8.405 | 118.534 |
| K94 | 7.581 | 117.212 |
| K95 | 7.660 | 116.793 |
| L96 | 8.540 | 120.770 |
| V97 | 8.814 | 119.657 |
| D98 | 8.330 | 117.849 |
| Q99 | 7.902 | 117.683 |
| I100 | 8.631 | 119.284 |
| E101 | 9.176 | 119.405 |
| F102 | 7.764 | 118.632 |
| Y103 | 7.985 | 120.937 |
| F104 | 7.491 | 112.011 |
| S105 | 7.721 | 117.776 |
| D106 | 9.260 | 123.776 |
| E107 | 8.479 | 116.583 |
| N108 | 7.340 | 115.049 |
| L109 | 8.726 | 118.996 |
| E110 | 7.249 | 113.843 |
| K111 | 7.013 | 114.564 |
| D112 | 7.841 | 123.120 |
| A113 | 7.898 | 126.203 |
| F114 | 8.430 | 119.168 |
| L115 | 7.633 | 122.265 |
| L116 | 8.610 | 120.427 |
| K117 | 8.091 | 116.584 |
| H118 | 7.227 | 117.321 |
| V119 | 7.875 | 120.947 |
| R120 | 8.031 | 113.038 |
| R121 | 7.635 | 117.990 |
| N122 | 7.089 | 116.031 |
| K123 | 9.110 | 126.062 |
| L124 | 7.959 | 117.451 |
| G125 | 8.151 | 109.937 |
| Y126 | 7.485 | 116.394 |
| V127 | 9.430 | 119.442 |
| S128 | 8.610 | 119.285 |
| V129 | 8.687 | 129.318 |
| K130 | 8.318 | 125.109 |
| L131 | 7.228 | 123.004 |

| Residue | δ($^1$H) [ppm] | δ($^{15}$N) [ppm] |
|---|---|---|
| L132 | 7.110 | 117.899 |
| T133 | 7.363 | 112.156 |
| S134 | 7.388 | 113.452 |
| F135 | 7.465 | 123.956 |
| K136 | 8.795 | 121.613 |
| K137 | - | - |
| V138 | 7.650 | 118.108 |
| K139 | 8.151 | 121.031 |
| H140 | 7.282 | 113.376 |
| L141 | 8.280 | 120.004 |
| T142 | 8.424 | 113.631 |
| R143 | 8.098 | 119.317 |
| D144 | 8.568 | 123.346 |
| W145 | 9.046 | 128.001 |
| R146 | 7.428 | 120.589 |
| T147 | 7.804 | 120.623 |
| T148 | 7.559 | 119.201 |
| A149 | 8.735 | 120.816 |
| H150 | 7.786 | 115.723 |
| A151 | 8.200 | 117.633 |
| L152 | 7.811 | 112.447 |
| K153 | 7.475 | 120.086 |
| Y154 | 8.041 | 115.658 |
| S155 | 7.101 | 113.626 |
| V156 | 9.186 | 121.890 |
| V157 | 7.516 | 116.785 |
| L158 | 7.545 | 120.866 |
| E159 | 8.969 | 119.155 |
| L160 | 8.503 | 125.624 |
| N161 | 8.385 | 120.089 |
| E162 | 8.631 | 118.234 |
| D163 | 7.249 | 113.843 |
| H164 | 7.725 | 111.541 |
| R165 | 8.139 | 112.310 |
| K166 | 8.136 | 118.474 |
| V167 | 9.204 | 119.294 |
| R168 | 8.790 | 123.190 |
| R169 | 9.104 | 120.880 |
| T170 | 8.050 | 114.175 |
| T171 | 7.540 | 115.556 |
| P172 | - | - |
| V173 | 8.645 | 125.913 |
| P174 | - | - |
| L175 | 7.697 | 117.844 |
| F176 | 8.878 | 125.372 |
| P177 | - | - |
| N178 | 8.614 | 114.897 |
| E179 | 8.040 | 120.152 |
| N180 | 8.281 | 118.671 |
| L181 | 8.140 | 123.725 |
| P182 | - | - |
| S183 | 8.014 | 121.981 |

**Table S6:** Structural statistics

| | |
|---|---|
| **Distance restraints** | |
| Total | 4116 |
| Intra-residue (\|i-j\|=0) | 520 |
| Sequential (\|i-j\|=1) | 949 |
| Medium-range (1<\|i-j\|<5) | 1219 |
| Long-range (\|i-j\|>=5) | 1428 |
| **Dihedral-angle restraints** | 154 |
| **Structural statistics** | |
| Distance restraints | 0.0029 ± 0.0005 Å |
| Dihedral angle restraints | 0.019 ± 0.004° |
| Max. distance restraint violation | 0.27 Å |
| Max. dihedral angle violation | 0.28° |
| Ramachandran Plot | |
| Most favored regions | 86.6 % |
| Additional allowed regions | 13.4 % |
| Generously allowed regions | 0.0 % |
| Disallowed regions | 0.0 % |
| Average RMSD (residues 78-183) | |
| Backbone atoms | 1.69 ± 0.34 Å |
| Heavy atoms | 1.76 ± 0.28 Å |
| Average RMSD (residues 85-176) | |
| Backbone atoms | 0.21 ± 0.04 Å |
| Heavy atoms | 0.62 ± 0.05 Å |

**Table S7:** Relaxation data of the 1:1 complex of HsLARP6(79-183) and A2M5 RNA in at 298 K in buffer containing 10 mM MES pH 6.5, 50 mM KCl, 10% $D_2O$, and 0.01 mg/mL DSS

| Residue | R1 | R1_ERR | R2 | R2_ERR | hetNOE | hetNOE_ERR | TAU_C [ns] | MW kDa | J0 [ns] | J0_ERR [ns] | JwN [ns] | JwN_ERR [ns] |
|---|---|---|---|---|---|---|---|---|---|---|---|---|
| G78 | | | | | | | | | | | | |
| E79 | #N/A | #N/A | #N/A | #N/A | #N/A | #N/A | | | | | | |
| D80 | 1.110 | 0.016 | 2.411 | 0.149 | -0.580 | 0.058 | 2.755 | 4.504 | 0.469 | 0.040 | 0.154 | 0.003 |
| L81 | 1.224 | 0.005 | 2.607 | 0.091 | -0.319 | 0.032 | 2.697 | 4.408 | 0.508 | 0.024 | 0.178 | 0.001 |
| E82 | 1.327 | 0.006 | 3.361 | 0.059 | -0.213 | 0.021 | 3.212 | 5.251 | 0.697 | 0.016 | 0.196 | 0.001 |
| Q83 | 1.308 | 0.010 | 4.699 | 0.120 | -0.027 | 0.003 | 4.280 | 6.997 | 1.062 | 0.032 | 0.200 | 0.002 |
| E84 | 1.277 | 0.013 | 6.766 | 0.171 | 0.202 | 0.020 | 5.585 | 9.131 | 1.625 | 0.046 | 0.202 | 0.002 |
| W85 | 1.024 | 0.008 | 11.231 | 0.211 | 0.252 | 0.025 | 8.599 | 14.059 | 2.859 | 0.055 | 0.164 | 0.001 |
| K86 | 0.776 | 0.010 | 16.297 | 0.699 | 0.431 | 0.043 | 12.232 | 19.998 | 4.255 | 0.183 | 0.127 | 0.002 |
| P87 | #N/A | #N/A | #N/A | #N/A | #N/A | #N/A | | | | | | |
| P88 | #N/A | #N/A | #N/A | #N/A | #N/A | #N/A | | | | | | |
| D89 | 0.647 | 0.006 | 18.454 | 0.250 | 0.724 | 0.072 | 14.371 | 23.494 | 4.855 | 0.066 | 0.111 | 0.002 |
| E90 | 0.629 | 0.009 | 19.869 | 0.430 | 0.822 | 0.082 | 15.148 | 24.766 | 5.237 | 0.120 | 0.110 | 0.002 |
| E91 | 0.654 | 0.011 | 20.700 | 0.229 | 0.777 | 0.078 | 15.168 | 24.797 | 5.456 | 0.063 | 0.113 | 0.002 |
| L92 | 0.662 | 0.012 | 22.847 | 0.600 | 0.750 | 0.075 | 15.866 | 25.939 | 6.030 | 0.161 | 0.114 | 0.002 |
| I93 | 0.589 | 0.025 | 23.196 | 1.308 | 0.799 | 0.080 | 16.989 | 27.775 | 6.134 | 0.341 | 0.102 | 0.004 |
| K94 | 0.635 | 0.015 | 23.223 | 0.424 | 0.808 | 0.081 | 16.343 | 26.718 | 6.135 | 0.116 | 0.110 | 0.003 |
| K95 | 0.641 | 0.016 | 22.660 | 0.626 | 0.829 | 0.083 | 16.068 | 26.268 | 5.984 | 0.168 | 0.112 | 0.003 |
| L96 | 0.663 | 0.016 | 21.377 | 0.376 | 0.748 | 0.075 | 15.313 | 25.035 | 5.636 | 0.099 | 0.114 | 0.003 |
| V97 | 0.636 | 0.015 | 22.129 | 0.632 | 0.888 | 0.089 | 15.929 | 26.042 | 5.842 | 0.176 | 0.112 | 0.003 |
| D98 | 0.673 | 0.015 | 23.137 | 0.533 | 0.821 | 0.082 | 15.828 | 25.876 | 6.107 | 0.143 | 0.117 | 0.003 |
| Q99 | 0.663 | 0.012 | 22.012 | 0.474 | 0.782 | 0.078 | 15.547 | 25.418 | 5.806 | 0.129 | 0.115 | 0.002 |
| I100 | #N/A | #N/A | #N/A | #N/A | #N/A | #N/A | | | | | | |
| E101 | 0.639 | 0.020 | 23.386 | 0.574 | 0.849 | 0.085 | 16.359 | 26.744 | 6.179 | 0.152 | 0.112 | 0.004 |
| F102 | 0.606 | 0.009 | 24.195 | 0.555 | 0.787 | 0.079 | 17.108 | 27.969 | 6.399 | 0.144 | 0.105 | 0.002 |
| Y103 | 0.634 | 0.014 | 24.716 | 0.635 | 0.900 | 0.090 | 16.898 | 27.626 | 6.536 | 0.174 | 0.112 | 0.003 |
| F104 | 0.628 | 0.018 | 21.404 | 0.568 | 0.841 | 0.084 | 15.765 | 25.774 | 5.649 | 0.150 | 0.110 | 0.003 |
| S105 | 0.635 | 0.009 | 24.184 | 0.510 | 0.827 | 0.083 | 16.699 | 27.300 | 6.393 | 0.132 | 0.111 | 0.002 |
| D106 | 0.653 | 0.018 | 20.442 | 0.464 | 0.808 | 0.081 | 15.085 | 24.662 | 5.387 | 0.121 | 0.113 | 0.003 |
| E107 | 0.686 | 0.013 | 19.639 | 0.193 | 0.766 | 0.077 | 14.398 | 23.539 | 5.167 | 0.050 | 0.118 | 0.003 |
| N108 | 0.671 | 0.013 | 19.117 | 0.504 | 0.845 | 0.084 | 14.359 | 23.475 | 5.030 | 0.133 | 0.117 | 0.002 |
| L109 | 0.684 | 0.022 | 22.427 | 0.518 | 0.826 | 0.083 | 15.454 | 25.265 | 5.915 | 0.139 | 0.119 | 0.004 |
| E110 | #N/A | #N/A | #N/A | #N/A | #N/A | #N/A | | | | | | |
| K111 | 0.657 | 0.008 | 19.440 | 0.423 | 0.712 | 0.071 | 14.651 | 23.952 | 5.117 | 0.110 | 0.112 | 0.002 |
| D112 | 0.593 | 0.014 | 17.379 | 0.384 | 0.621 | 0.062 | 14.572 | 23.823 | 4.573 | 0.104 | 0.100 | 0.003 |
| A113 | 0.663 | 0.011 | 19.802 | 0.529 | 0.741 | 0.074 | 14.716 | 24.059 | 5.214 | 0.140 | 0.114 | 0.002 |
| F114 | 0.678 | 0.022 | 23.596 | 0.579 | 0.728 | 0.073 | 15.939 | 26.058 | 6.228 | 0.154 | 0.116 | 0.004 |
| L115 | 0.746 | 0.021 | 22.346 | 0.569 | 0.821 | 0.082 | 14.743 | 24.103 | 5.885 | 0.157 | 0.130 | 0.004 |
| L116 | 0.694 | 0.020 | 21.925 | 1.048 | 0.827 | 0.083 | 15.149 | 24.766 | 5.779 | 0.278 | 0.121 | 0.004 |
| K117 | 0.671 | 0.019 | 21.277 | 0.371 | 0.733 | 0.073 | 15.186 | 24.827 | 5.608 | 0.099 | 0.115 | 0.003 |
| H118 | 0.682 | 0.014 | 21.716 | 0.811 | 0.769 | 0.077 | 15.219 | 24.881 | 5.724 | 0.212 | 0.118 | 0.003 |
| V119 | 0.658 | 0.021 | 21.901 | 1.367 | 0.674 | 0.067 | 15.565 | 25.447 | 5.776 | 0.373 | 0.112 | 0.004 |
| R120 | 0.633 | 0.026 | 18.854 | 0.956 | 0.705 | 0.070 | 14.698 | 24.029 | 4.964 | 0.251 | 0.108 | 0.005 |
| R121 | 0.654 | 0.015 | 22.563 | 0.601 | 0.703 | 0.070 | 15.866 | 25.939 | 5.955 | 0.163 | 0.112 | 0.003 |
| N122 | 0.705 | 0.014 | 22.109 | 0.875 | 0.833 | 0.083 | 15.093 | 24.676 | 5.827 | 0.235 | 0.123 | 0.003 |
| K123 | 0.747 | 0.011 | 19.459 | 0.417 | 0.727 | 0.073 | 13.706 | 22.407 | 5.110 | 0.116 | 0.128 | 0.002 |
| L124 | 0.680 | 0.049 | 27.685 | 2.583 | 0.711 | 0.071 | 17.279 | 28.249 | 7.323 | 0.675 | 0.116 | 0.008 |
| G125 | 0.686 | 0.020 | 25.445 | 3.075 | 0.801 | 0.080 | 16.462 | 26.913 | 6.723 | 0.839 | 0.119 | 0.004 |
| Y126 | 0.643 | 0.021 | 22.099 | 0.708 | 0.980 | 0.098 | 15.834 | 25.887 | 5.835 | 0.186 | 0.114 | 0.004 |
| V127 | 0.646 | 0.019 | 20.636 | 1.158 | 0.762 | 0.076 | 15.236 | 24.908 | 5.440 | 0.323 | 0.112 | 0.003 |
| S128 | #N/A | #N/A | #N/A | #N/A | #N/A | #N/A | | | | | | |
| V129 | 0.828 | 0.048 | 24.624 | 2.165 | 0.770 | 0.077 | 14.687 | 24.011 | 6.483 | 0.578 | 0.143 | 0.009 |
| K130 | 0.666 | 0.018 | 18.657 | 0.787 | 0.934 | 0.093 | 14.237 | 23.275 | 4.909 | 0.216 | 0.118 | 0.004 |
| L131 | 0.690 | 0.019 | 22.482 | 1.107 | 0.813 | 0.081 | 15.397 | 25.172 | 5.929 | 0.292 | 0.120 | 0.004 |
| L132 | 0.676 | 0.022 | 21.734 | 0.983 | 0.856 | 0.086 | 15.291 | 24.998 | 5.731 | 0.264 | 0.118 | 0.004 |
| T133 | 0.657 | 0.019 | 19.279 | 0.676 | 0.737 | 0.074 | 14.578 | 23.832 | 5.074 | 0.181 | 0.113 | 0.003 |
| S134 | 0.681 | 0.014 | 21.872 | 0.665 | 0.831 | 0.083 | 15.282 | 24.983 | 5.767 | 0.182 | 0.119 | 0.003 |
| F135 | 0.740 | 0.014 | 21.749 | 0.705 | 0.885 | 0.088 | 14.599 | 23.867 | 5.727 | 0.194 | 0.130 | 0.003 |
| K136 | 0.692 | 0.017 | 25.132 | 0.796 | 0.830 | 0.083 | 16.287 | 26.628 | 6.639 | 0.205 | 0.121 | 0.003 |
| K137 | #N/A | #N/A | #N/A | #N/A | #N/A | #N/A | | | | | | |
| V138 | #N/A | #N/A | #N/A | #N/A | #N/A | #N/A | | | | | | |
| K139 | 0.687 | 0.046 | 53.562 | 6.139 | 0.899 | 0.090 | 24.079 | 39.365 | 14.257 | 1.651 | 0.121 | 0.008 |
| H140 | 0.756 | 0.015 | 25.602 | 0.708 | 0.707 | 0.071 | 15.705 | 25.675 | 6.754 | 0.184 | 0.129 | 0.003 |
| L141 | 0.662 | 0.016 | 24.155 | 1.260 | 0.794 | 0.079 | 16.324 | 26.687 | 6.381 | 0.347 | 0.115 | 0.003 |

| Residue | R1 | R1_ERR | R2 | R2_ERR | hetNOE | hetNOE_ERR | TAU_C [ns] | MW kDa | J0 [ns] | J0_ERR [ns] | JwN [ns] | JwN_ERR [ns] |
|---|---|---|---|---|---|---|---|---|---|---|---|---|
| T142 | 0.618 | 0.013 | 20.044 | 0.482 | 0.752 | 0.075 | 15.361 | 25.113 | 5.285 | 0.127 | 0.107 | 0.003 |
| R143 | 0.610 | 0.020 | 21.487 | 1.094 | 0.639 | 0.064 | 16.031 | 26.208 | 5.672 | 0.296 | 0.103 | 0.004 |
| D144 | 0.664 | 0.017 | 19.433 | 0.921 | 0.805 | 0.081 | 14.568 | 23.816 | 5.115 | 0.252 | 0.115 | 0.003 |
| W145 | 0.717 | 0.027 | 20.674 | 2.445 | 0.769 | 0.077 | 14.452 | 23.627 | 5.440 | 0.656 | 0.124 | 0.005 |
| R146 | 0.720 | 0.016 | 24.474 | 1.192 | 0.867 | 0.087 | 15.740 | 25.733 | 6.459 | 0.314 | 0.126 | 0.003 |
| T147 | 0.629 | 0.019 | 23.901 | 0.845 | 0.841 | 0.084 | 16.679 | 27.268 | 6.318 | 0.221 | 0.110 | 0.004 |
| T148 | 0.712 | 0.011 | 23.240 | 0.567 | 0.791 | 0.079 | 15.415 | 25.202 | 6.129 | 0.150 | 0.123 | 0.002 |
| A149 | 0.729 | 0.017 | 22.696 | 0.938 | 0.818 | 0.082 | 15.035 | 24.580 | 5.981 | 0.251 | 0.127 | 0.003 |
| H150 | 0.629 | 0.018 | 20.751 | 0.659 | 0.820 | 0.082 | 15.494 | 25.330 | 5.474 | 0.173 | 0.110 | 0.003 |
| A151 | 0.678 | 0.020 | 21.683 | 0.329 | 0.827 | 0.083 | 15.245 | 24.923 | 5.716 | 0.087 | 0.118 | 0.004 |
| L152 | 0.662 | 0.018 | 19.142 | 0.854 | 0.821 | 0.082 | 14.474 | 23.663 | 5.038 | 0.227 | 0.115 | 0.003 |
| K153 | 0.681 | 0.014 | 23.364 | 0.579 | 0.806 | 0.081 | 15.813 | 25.853 | 6.167 | 0.152 | 0.118 | 0.003 |
| Y154 | 0.674 | 0.011 | 21.209 | 0.621 | 0.799 | 0.080 | 15.124 | 24.726 | 5.590 | 0.166 | 0.117 | 0.002 |
| S155 | 0.636 | 0.009 | 16.900 | 0.225 | 0.776 | 0.078 | 13.851 | 22.644 | 4.441 | 0.060 | 0.110 | 0.002 |
| V156 | 0.659 | 0.012 | 17.828 | 0.537 | 0.776 | 0.078 | 13.980 | 22.856 | 4.686 | 0.149 | 0.114 | 0.003 |
| V157 | 0.727 | 0.022 | 19.865 | 0.777 | 0.734 | 0.073 | 14.053 | 22.975 | 5.222 | 0.206 | 0.125 | 0.004 |
| L158 | 0.682 | 0.027 | 22.351 | 0.859 | 0.679 | 0.068 | 15.443 | 25.248 | 5.894 | 0.234 | 0.116 | 0.005 |
| E159 | 0.640 | 0.015 | 21.281 | 0.729 | 0.738 | 0.074 | 15.564 | 25.445 | 5.613 | 0.195 | 0.110 | 0.003 |
| L160 | 0.626 | 0.013 | 17.671 | 0.397 | 0.697 | 0.070 | 14.289 | 23.361 | 4.648 | 0.106 | 0.107 | 0.003 |
| N161 | 0.655 | 0.018 | 20.717 | 0.837 | 0.859 | 0.086 | 15.164 | 24.791 | 5.461 | 0.226 | 0.115 | 0.003 |
| E162 | 0.678 | 0.027 | 22.640 | 0.815 | 0.761 | 0.076 | 15.601 | 25.506 | 5.972 | 0.208 | 0.117 | 0.005 |
| D163 | #N/A | #N/A | #N/A | #N/A | #N/A | #N/A | | | | | | |
| H164 | 0.650 | 0.019 | 21.436 | 0.446 | 0.825 | 0.082 | 15.493 | 25.328 | 5.654 | 0.121 | 0.113 | 0.004 |
| R165 | 0.689 | 0.028 | 20.938 | 0.802 | 0.706 | 0.071 | 14.854 | 24.285 | 5.514 | 0.216 | 0.118 | 0.005 |
| K166 | 0.713 | 0.018 | 21.450 | 0.598 | 0.764 | 0.076 | 14.776 | 24.157 | 5.649 | 0.162 | 0.123 | 0.004 |
| V167 | 0.736 | 0.031 | 21.566 | 0.972 | 0.821 | 0.082 | 14.574 | 23.826 | 5.677 | 0.265 | 0.128 | 0.006 |
| R168 | 0.696 | 0.033 | 25.088 | 0.894 | 0.900 | 0.090 | 16.220 | 26.518 | 6.627 | 0.239 | 0.123 | 0.006 |
| R169 | 0.710 | 0.020 | 25.394 | 1.206 | 0.902 | 0.090 | 16.163 | 26.424 | 6.707 | 0.322 | 0.125 | 0.004 |
| T170 | 0.678 | 0.014 | 20.467 | 0.431 | 0.704 | 0.070 | 14.800 | 24.196 | 5.389 | 0.120 | 0.116 | 0.003 |
| T171 | 0.704 | 0.009 | 18.868 | 0.242 | 0.716 | 0.072 | 13.907 | 22.736 | 4.958 | 0.064 | 0.121 | 0.002 |
| P172 | #N/A | #N/A | #N/A | #N/A | #N/A | #N/A | | | | | | |
| V173 | 0.702 | 0.014 | 15.274 | 0.250 | 0.575 | 0.057 | 12.469 | 20.385 | 3.993 | 0.069 | 0.118 | 0.003 |
| P174 | #N/A | #N/A | #N/A | #N/A | #N/A | #N/A | | | | | | |
| L175 | 0.697 | 0.010 | 17.709 | 0.196 | 0.515 | 0.052 | 13.527 | 22.115 | 4.646 | 0.053 | 0.116 | 0.002 |
| F176 | 0.770 | 0.015 | 15.085 | 0.234 | 0.533 | 0.053 | 11.789 | 19.273 | 3.933 | 0.063 | 0.128 | 0.003 |
| P177 | #N/A | #N/A | #N/A | #N/A | #N/A | #N/A | | | | | | |
| N178 | 0.855 | 0.014 | 15.251 | 0.284 | 0.391 | 0.039 | 11.219 | 18.342 | 3.963 | 0.074 | 0.139 | 0.002 |
| E179 | 1.042 | 0.005 | 11.876 | 0.114 | 0.290 | 0.029 | 8.785 | 14.363 | 3.030 | 0.029 | 0.167 | 0.001 |
| N180 | 1.221 | 0.011 | 6.667 | 0.084 | 0.135 | 0.014 | 5.694 | 9.308 | 1.605 | 0.021 | 0.191 | 0.002 |
| L181 | 1.160 | 0.004 | 3.353 | 0.078 | -0.337 | 0.034 | 3.608 | 5.899 | 0.718 | 0.021 | 0.168 | 0.001 |
| P182 | #N/A | #N/A | #N/A | #N/A | #N/A | #N/A | | | | | | |
| S183 | 0.856 | 0.002 | 1.497 | 0.017 | -0.799 | 0.080 | 2.096 | 3.427 | 0.261 | 0.005 | 0.114 | 0.002 |

**Table S8:** $K_D$ and ΔΔG of A2M5 binding to wild type and mutant La domain variants

| Mutation | $K_D$ | $K_D$ error[a] | ΔΔG | ΔΔG Error[b] |
|---|---|---|---|---|
| WT | 5.55 | 0.19 | - | - |
| Q99A | 3.18 | 0.27 | -0.33 | 0.05 |
| F102A | 8.88 | 1.26 | 0.28 | 0.09 |
| K111A | 4.34 | 0.43 | -0.15 | 0.06 |
| F114A | 309.37 | 49.67 | 2.38 | 0.10 |
| K117A | 21.46 | 1.12 | 0.80 | 0.04 |
| H118A | 19.08 | 1.35 | 0.73 | 0.05 |
| R121A | 43.54 | 2.92 | 1.22 | 0.04 |
| N122A | 18.74 | 0.56 | 0.72 | 0.03 |
| K123A | 28.25 | 1.89 | 0.96 | 0.04 |
| K130A | 62.46 | 7.51 | 1.43 | 0.07 |
| K136A | 9.71 | 0.29 | 0.33 | 0.03 |
| K137A | 2.69 | 0.39 | -0.43 | 0.09 |
| K139A | 36.92 | 0.83 | 1.12 | 0.02 |
| H140A | 12.50 | 0.55 | 0.48 | 0.03 |
| R165A | 15.84 | 1.32 | 0.62 | 0.05 |
| K166A | 203.72 | 20.60 | 2.13 | 0.06 |

[a] Standard deviation in $K_D$ from fit of MST data sets with 3 replicates

[b] Standard deviation calculated by error propagation
